# Multimodal synapse analysis reveals limitations in transplanted neuron integration mediated by TREM2

**DOI:** 10.1101/2025.01.31.635250

**Authors:** Yvette Zarb, Oskar Markkula, Eva-Maria Schentarra, Manja Thorwirth, Marfa Fernanda Martfnez-Reza, Emanuele Paoli, Maria L. Richter, Mihail Todorov, Christina Koupourtidou, Veronika Schwarz, Georg Kislinger, Alexandra Mezydlo, Chu Lan Lao, Matteo Puglisi, Michael Willem, Jovica Ninkovic, Karl-Klaus Conzelmann, Martina Schifferer, Christian Haass, Martin Kerschensteiner, Ruben Portugues, Ralf Jungmann, Conny Kopp-Scheinpflug, Magdalena Gotz

## Abstract

Neuron transplantation offers a promising approach for circuit restoration after injury, but depends crucially on adequate synaptic integration. Yet, our knowledge of the effects of the host injury environment on transplanted neuron (tN) network integration remains limited. Here, we used a stab wound injury model to examine how tN synapses mature and integrate using multimodal read-outs. Morphological, ultrastructural and functional electrophysiological analysis of tN synapses indicated that tN integration and functional maturation are partially stunted. Spatial transcriptomics revealed persistent inflammatory signatures at the transplant site including *Trem2* upregulation. Strikingly, in an environment devoid of TREM2, synaptic integration as well as functional maturation of tNs, including action potential maturation and spontaneous firing, were almost fully rescued. Spatial transcriptomics further showed reduced inflammatory pathways and improved maturation of tNs in *Trem2^-/-^* brains. This study highlights the importance of tackling chronic inflammation for adequate tN integration.

## Introduction

Neurons lost due to brain injury or neurodegenerative disease can so far not be regenerated, except by transplantation of new neurons {*1*}. The aim is to restore neural circuit function, requiring adequate neuronal integration. In a proof-of-concept study using a neuron ablation model causing little inflammation, transplanted neurons (tNs) displayed adequate circuit integration, such as area-appropriate brain-wide input connectome and correct receptive field properties {*2*}. Likewise, tNs extend processes and connect to regions similar to their endogenous counterparts in different disease models {*3–6*}. Mapping the brain-wide input connectome to tNs by monosynaptic Rabies virus tracing (mRABV) showed synapse loss in the host circuitry as the main determinant of tN integration {*2*, *7*, *8*}. Yet, our knowledge about tN synaptic connectivity is still very limited as typically only one or two synaptic read-outs were used. For example, spines are often taken as a proxy for excitatory synapses, as normally the majority of spines have glutamate receptors and most excitatory synapses are on spines in the cerebral cortex {*9*}. However, it is not clear to what extent this is also the case for tNs. Moreover, monosynaptic Rabies virus tracing allows monitoring brain-wide neuronal connectivity, but to which extent these connections are functional on tNs after brain injury is not known. We therefore set out here to use a multimodal approach to analyze how the synaptic connectivity of tNs is refined on the structural, ultrastructural and functional levels.

Moreover, many studies investigated tN integration either in inflammatory-stunted environments, to allow for human neuron transplantation, or used of injury models that do not generate an immune response. However, the host injury condition plays a major role in tN integration. Aging, amyloid deposition and traumatic brain injury result in differences in the input connectome traced by RABV with proteomic hallmarks of inflammation correlating best to the differences in tN connectivity {*7*, *8*}. However, the role of inflammation in sculpting tN integration has not been directly explored. Microglia are one of the main contributors to neuroinflammation, and circuit remodeling in development {*10–13*} and disease {*14–17*}, but their contribution to tN integration is still poorly studied. Here we therefore used a multimodal synapse analysis to assess the integration of tNs in the stab wound injury, revealing restricted maturation and connectivity. Spatial transcriptomics detected higher persisting expression of activated microglia signatures around tNs including a *Trem2* signature. Transplanting in an environment devoid of TREM2, reduced the inflammation around tNs and resulted in a striking effect on improving neuronal differentiation and synaptic maturation as well as their brain-wide input connectome.

## Results

### tNs exhibit persistently elevated spine density

During development, synaptic connections of neurons are refined for maturity through the pruning of spines and the dendritic arbor {*18*}. Here, we first aimed to monitor if entire tN dendrites were pruned or reshaped over time. In our previous analysis of tNs after neuronal ablation, major changes in reshaping basal dendrites were restricted to the first weeks after transplantation {*2*}. We therefore performed longitudinal 2-photon live imaging from 1- and 3-months post transplantation (mpt) to examine individual dendrites of the tNs in a stab wound injury (SWI) condition (Fig. S1A). As before {*2*, *7*, *8*}, tNs were obtained from mouse embryonic day (E) 14 cerebral cortex, cultured *in vitro* for 3-5 days and infected with a retrovirus expressing CAG-GFP at 1 day *in vitro* (DIV). Cells were transplanted 1 week after SWI and imaging started at 1mpt until 3mpt (Fig. S1B). The dendritic processes from neurons that were equally well visible at both 1mpt and 3mpt were recorded. When comparing the dendritic length of tNs at 1mpt and 3mpt (Fig. S1C), no significant difference was detected (1mpt – 87 ± 50 µm; 3mpt – 92 ± 47 µm), indicating that no major dendritic pruning of tNs occurred between 1mpt and 3mpt. Thus, transplantation of neurons after SWI does not result in prolonged or exacerbated dendrite pruning. However, the absence of dendritic pruning does not reveal whether the synapses on these dendrites are maturing appropriately, prompting us to examine spine density as a proxy for synapse formation.

As an initial assessment of the maturation of tNs and synapse formation, we analyzed the spine density of GFP-labeled tNs at 1 and 3mpt. As control, spine density of endogenous neurons in the intact cortex were assessed by using a SAD-ΔG RABV expressing GFP (Fig. 1A), which preferentially infects neurons {*19*}. Notably, these neurons did not exhibit any morphological signs of RABV-mediated toxicity. For tNs, spine density was quantified from dendrites expressing GFP projecting outside the transplanted area (Fig. 1A; Fig. 1B). When compared to endogenous neurons labeled with RABV, a significant 2-fold increase in spine density of tNs was observed (Fig. 1C). No significant difference in spine density was observed for tNs between 1mpt and 3mpt (Fig. 1C).

**Figure 1.**
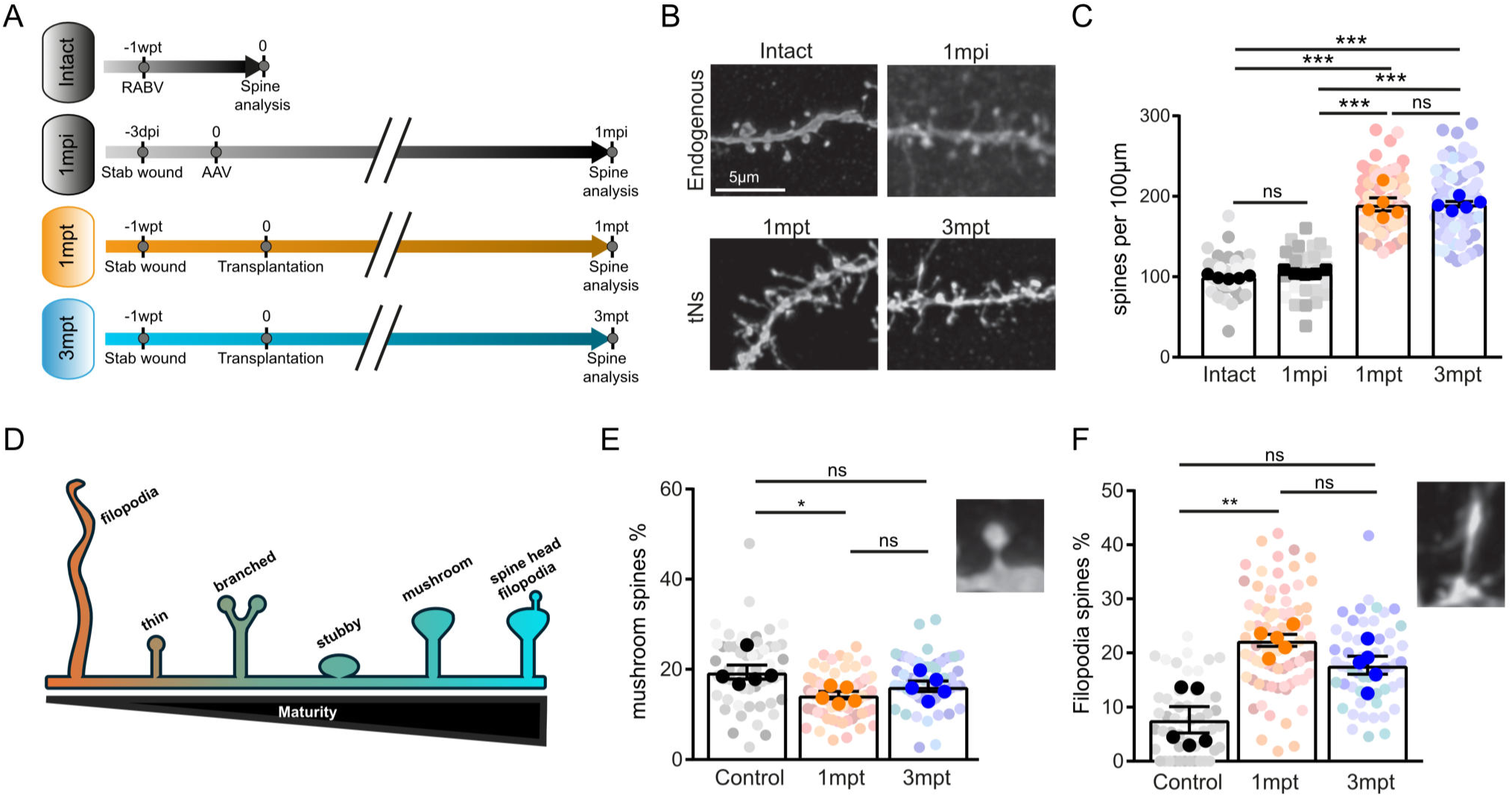
Dendritic tN spines mature between 1 and 3mpt. **(A)** Experimental setup used to analyze spine density and type. **(B)** Images of dendritic processes from endogenous pyramidal neurons in intact brain labelled with RABV (top left) and 1mpi labelled with AAVs (top right), and transplanted neurons at 1mpt (bottom left) and 3mpt (bottom right). **(C)** Spine density quantification of 1mpt, 3mpt and endogenous neurons (mice (n) = 5, dendrites = 20-77). **(D)** Illustration representing the different types of spine types. **(E, F)** Images and their respective spine type quantification in 1mpt, 3mpt and endogenous neurons in intact brain of **(E)** mushroom and **(F)** filopodia spine types. (mice (n) = 5, dendrites = 53-75). In graphs, solid colors represent per-animal means; lighter shades represent individual measurements, with distinct shades denoting each mouse. All data are presented as means ± SEM. All statistics were performed on the means of individual mice - ns p >0.05, * p <0.05, ** p <0.005, by Kruskal-Wallis test with Dunn’s test (C, E, F).

To explore if spine density is affected by the SWI, we also analyzed endogenous neurons after SWI only without transplantation. To eliminate any confounding effects of RABV, we labeled endogenous cortical cells with a cocktail of PHP.B-capsid AAVs: pAAV-gfaABC1D-iCre, pAAV-CBh-FLEX-EGFP, and pAAV-CBh-FLEX-DsRed2-W3SL, 3-days post injury. For this analysis, we took advantage of the leaky activity of this promoter for the sparse labeling of endogenous neurons. The mice were sacrificed at 1-month post injury (mpi; Fig. 1A). Notably, the spine density of endogenous neurons at 1mpi assessed as above did not significantly differ from the spine density of endogenous pyramidal neurons in intact brains, labeled with RABV (Fig. 1C), further indicating the lack of toxicity of our RABV constructs. These results suggest that the increased spine density of tNs at 1mpt and 3mpt after SWI, is a feature of tN development in the SWI environment rather than an adult neuronal response to injury.

Previous studies suggested that the culture period prior to transplantation does not significantly influence tN connectivity by using RABV {*2*, *7*, *8*}. Nevertheless, we aimed to control for a possible influence of the culture period prior to transplantation by transplanting acutely dissociated neurons from cerebral cortices of E18 β-*Actin-GFP* mice 1 week after SWI. Notably, these tNs had a spine density comparable to that of E14 cultured tNs (Fig. S1D, E). Taken together, tNs maintain a high spine density up to 3mpt, which may reflect a persistent degree of immaturity. To determine whether this elevated spine density reflects genuine synaptic maturation, we next analyzed the spine type composition as a readout of synapse maturity.

### Spines of tNs mature between 1 and 3mpt

Synaptic strength and transience have been linked to spine type {*20*}, as for example mushroom spines indicate more mature spines with mature/stable synapses in most cases ((*20*]; Fig. 1D). Consequently, we quantified the spine types of endogenous neurons and tNs labeled with GFP at 1 and 3mpt (Fig. 1D-F, Fig. S1F-G). Given the significant increase of tN spine density (Fig. 1C) resulting in elevated quantities of all spine types, we proceeded to compare the percentage of spine types in relation to the total spine density. Interestingly, the percentage of mushroom spines was significantly lower in tNs at 1mpt compared to endogenous neurons (Fig. 1E). However, at 3mpt the percentage of mushroom spines increased to levels comparable to endogenous neurons (Fig. 1E). These data indicate the maturation of tN spines over time.

We then analyzed the percentage of filopodial spines, which are typically associated with immaturity {*21*}. At 1mpt, the percentage of filopodial spines was significantly higher (almost 3-fold) on tNs compared to endogenous neurons (Fig. 1F). Despite the decrease in filopodia spines at 3mpt (17.8 ± 7.4%), tN dendrites still had an approximately 2-fold higher percentage of filopodia spines compared to endogenous neurons, albeit no longer significantly different (Fig. 1F). Other spine subtypes, such as branched (Fig. S1F), thin (Fig. S1G), spine head filopodia (Fig. S1H), and stubby spines (Fig. S1I) were also quantified, showing significant differences only in the stubby spines (significantly reduced in their proportion on tNs) and branched spines that were significantly increased on the tNs. While these spine types provide less direct information about spine maturity, stubby spines have been described as immature spines, less likely to be connected to a synapse{*22*}, and were found to be decreased in stress conditions {*23*}. Conversely, branched spines were found to be increased in tNs compared to endogenous neurons, in line with their appearance during synaptogenesis or plasticity conditions {*24*}. All-in-all these data indicate a significant degree of spine maturation between 1 and 3mpt, as well as some degree of immaturity persisting until 3mpt. While spine morphology provides an initial indication of synaptic status, it does not reveal whether these spines harbor functional presynaptic partners. We therefore turned to 3D ultrastructural analysis to examine the synaptic composition of tN dendrites directly.

### 3D ultrastructural analysis of tNs shows many empty spines and synapses on dendritic shafts

To identify the location and opposition of the pre- and postsynaptic components of tN synapses, we analyzed their three-dimensional (3D) ultrastructure using automated tape-collecting ultramicrotome scanning electron microscopy (ATUM-SEM; Fig. S2A). For this analysis, we segmented dendrites from tNs at 1mpt and 3mpt and as control, we used the published 3D-EM dataset of Layer II/III adult mouse visual cortex {*25*}. GFP-positive tNs were identified by the dark spots in the soma or dendritic stretches following immunogold labelling (arrowheads in Fig. S2B), whereas the control neurons were identified based on morphology. Control immunogold-labeling of GFP in brain tissue of mice expressing the fluorescent protein either Cux2-positive cells after Cre-mediated recombination (Cux2-CreERT2 x GFPrep) (Fig. S2C) and under the Cx3cr1 promoter (*Cx3cr1^GFP^* ; Fig. S2D) confirmed the reliability of our immunostaining by labelling upper layer neurons and microglia, respectively.

Using this experimental paradigm, we analyzed the synaptic connectivity of GFP-labelled tNs by segmenting the 3D ultrastructure of a selection of tN and control dendrites, together with the observed synapses (Fig. 2A). In the dataset, we annotated synapses at dendritic spines (Fig. S2E magenta arrow) and shafts (Fig. S2E blue arrow), and also spines lacking a presynaptic partner (Fig. S2E green arrow). Quantifying the synaptic structures on spines in the three conditions revealed a surprisingly high proportion, almost half at 1mpt, of tN spines without an associated synapse (empty spines), while this was much less frequent (13.4 ± 3.1%) in the control neurons (Fig. 2B). This proportion decreases at 3mpt, but not to control levels. Unexpectedly, spine synapses only make up less than a third of all synapses on tNs at 1mpt with the vast majority of the synapses on tNs on the shaft (Fig. 2C). The number of spine synapses doubled at 3mpt, albeit not yet reaching control levels (Fig. 2C). Interestingly, both at the innervation (Fig. 2B) and at the synapse location (Fig. 2C) level, the 3mpt quantifications indicate partial restoration to control levels, suggesting an incomplete maturation trajectory of tNs between 1mpt and 3mpt.

**Figure 2.**
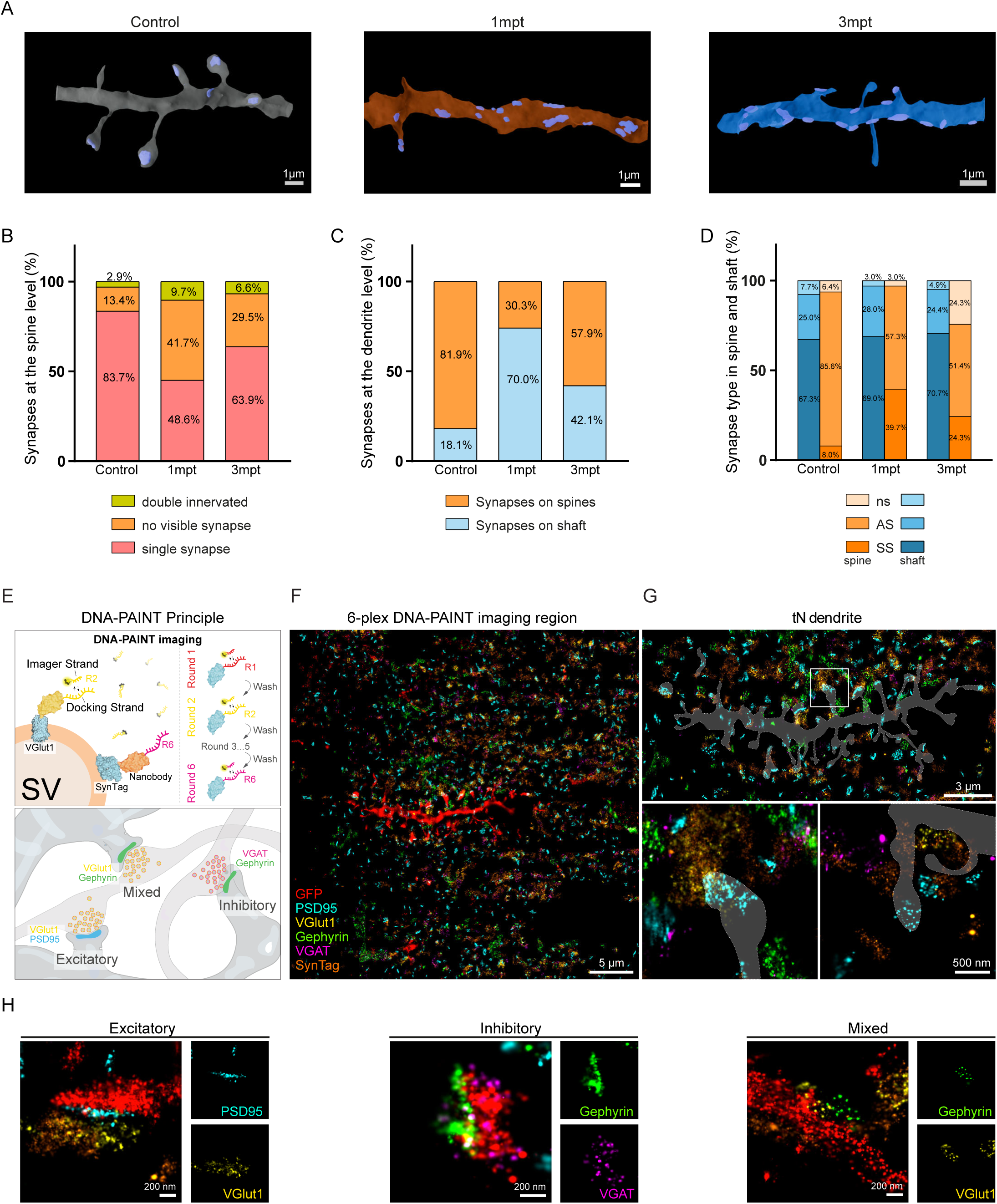
Synapses on tNs show immature hallmarks at 1mpt and 3mpt. **(A)** Reconstructions of dendrites in control (left), 1mpt (middle), and 3mpt (right), and respective synapse analysis **(B-C)**. Percentage of synapses on spines **(B)**, and the overall dendrite **(C)**. **(D)** Percentage of asymmetrical synapses and symmetrical synapses distribution in spines (orange) and shaft (blue) (3 mice (1 per condition), dendrites = 5-17). **(E)** DNA-PAINT principal and general experimental design. In DNA-PAINT, fluorescently labeled imager strands transiently bind to complementary docking strands attached to the target protein via immunolabeling, enabling single-molecule localization microscopy (SMLM) imaging and visualization of multiple synaptic proteins within the same sample. **(F)** Six-plex DNA-PAINT imaging region the antigens as indicated. **(G)** Inset of a tN dendrite from (F) and zoom-ins of spines with chemical synapses (bottom left), and empty spines lacking a presynapse (bottom right). **(H)** Example images of an excitatory (left), inhibitory (middle), and mixed (right) synapse.

To understand if these observations were due to data points skewing the data, we plotted the values of the different dendritic values for each condition and observed a small SEM in control and 3mpt conditions (Fig. S2F, G). The 1mpt condition was more variable, but when we compared the dendritic values according to tNs analyzed, there was no significant difference between the different neurons (Fig. S2H, I).

To further characterize the nature of synapses on tNs, we assessed if the synapses observed were symmetrical or asymmetrical (Fig. S2J). Symmetrical synapses typically are inhibitory, whereas asymmetrical synapses with a thick postsynaptic density (Fig. S2J cyan arrow), are excitatory {*26*}. As expected, the majority of synapses on spines in the control were asymmetrical synapses (85.6%; Fig. 2D), with most of the symmetrical synapses located on doubly innervated spines. However, this was not the case for tNs, where barely half of the spine synapses were asymmetrical (Fig. 2D). Conversely, shaft synapses were predominantly symmetrical in all three conditions (Fig. 2D). In summary, tNs in the SWI differ from adult pyramidal neurons in the cortex by a high fraction of spines without a synaptic partner (Fig. 2B) and a high proportion of shaft synapses (Fig. 2C), although this is partially improved at 3mpt. These results add to our previous results that tNs are more immature than endogenous neurons even at 3mpt, as a high degree of shaft synapses is reminiscent of immature neurons during development [*21*).

### Multiplexed high resolution synapse staining reveals many empty tN postsynapses at 3mpt

To assess tN synapses beyond the morphological level, we immunostained for pre- and postsynaptic proteins characteristic of inhibitory and excitatory chemical synapses on tNs. To achieve better resolution than possible with conventional immunofluorescence imaging techniques, we used the single molecule localization microscopy technique, DNA-PAINT, to image several synaptic protein targets simultaneously in nanoscale topography. DNA-PAINT has previously been used to resolve multiple synaptic proteins at super-resolution {*27*}. With this technique we were able to resolve the presynaptic vesicle pool and postsynaptic scaffold proteins of inhibitory and excitatory synapses in tissue simultaneously at a resolution of ∼20 nm (Fig. 2E, F). A synapse containing vGLUT1 and PSD95 was defined as excitatory and one containing gephyrin and vGAT as inhibitory (Fig. 2E, H). In addition, in line with previous data [*27*) we also observed the “mixed” synapses as previously identified by a postsynaptic gephyrin and presynaptic vGLUT1 signal (Fig. 2E, H). DNA-PAINT imaging revealed spines of tNs with excitatory synapses (Fig. 2G bottom left), but many spines were “empty” with only the PSD95 scaffold present devoid of any presynaptic vesicle pool protein (Fig. 2G bottom right), referred as PSD95 empty. We observed that full synapses, i.e. pre- and postsynaptic staining, consisted of the minority of the total synapses detected (n=180), with only 31.6% of excitatory synapses and 15.4% of inhibitory synapses, were full synapses. This aligned with our EM observation that tNs have a high proportion of “empty” spines, i.e. spines without a presynaptic partner. While these are distinct measurements, both indicate spines that lack a complete, functional synapse and are therefore consistent with the broader pattern of incomplete synaptic maturation of tNs. Further analysis revealed that around 4% of all the synapses were classified as mixed, which have previously been suggested to be immature {*27*}. All-in-all these results give us a snapshot of the synaptome architecture of tNs at 3mpt at unprecedented detail, hinting at an immature state persisting at 3mpt, highlighting the need to characterize the functional state of the tN synaptic connectivity. Having established that tN synapses are structurally and molecularly immature, we next asked whether this immaturity is also reflected at the functional level.

### tNs can fire action potentials but remain largely silent

To investigate the functional integration and maturation of tNs in the SWI, we performed whole-cell patch clamp recordings from acute slices of the cerebral cortices containing tNs at 1 or 3mpt. As controls, neighboring endogenous neurons adjacent to the area of transplantation were patched. To investigate the intrinsic maturation of tNs, we first focused on the analysis of action potentials. Upon depolarization, tNs were able to generate action potentials at both 1 and 3mpt, but with an increased half-width when compared to endogenous neurons in the same SW-injured brain region (Fig. 3A-D). While all neurons in the three conditions were able to generate increasing numbers of action potentials in response to depolarizing current injections (Fig. 3E-G), the frequency of the firing rate was lower for tNs, when compared to endogenous neurons (Fig. 3H). Moreover, we found an almost complete lack of spontaneous action potentials in tNs at 1mpt (Fig. 3J) with only a slight increase at 3mpt (Fig. 3K), which was very different from the neighboring endogenous neurons around the transplantation area in the same injured environment (Fig. 3I). This pattern was reflected in the high number of silent tNs, at 1mpt 80% (Fig. S3A middle) that slightly recovered to 60% at 3mpt (Fig. S3A right), still much different from endogenous neurons in the same area with only a quarter [24%) silent. A Pearson Chi-squared test of independence was performed on a 3x2 contingency table and resulted in statistically significant (p ≤ 0.01) association between treatment group (control, 1mpt, 3mpt) and activity state (silent vs active). Cumulative distributions of overall spontaneous activity show lower firing rates in tNs at both 1mpt and 3mpt compared to endogenous neurons (Fig. 3L). Input resistance (Fig. S3B), membrane time constant (Fig. S3C), and resting membrane potential (Fig. S3D), were not significantly different between the three groups. Although tNs are capable of firing action potentials, their half-width and firing frequency indicate an immature state.

**Figure 3.**
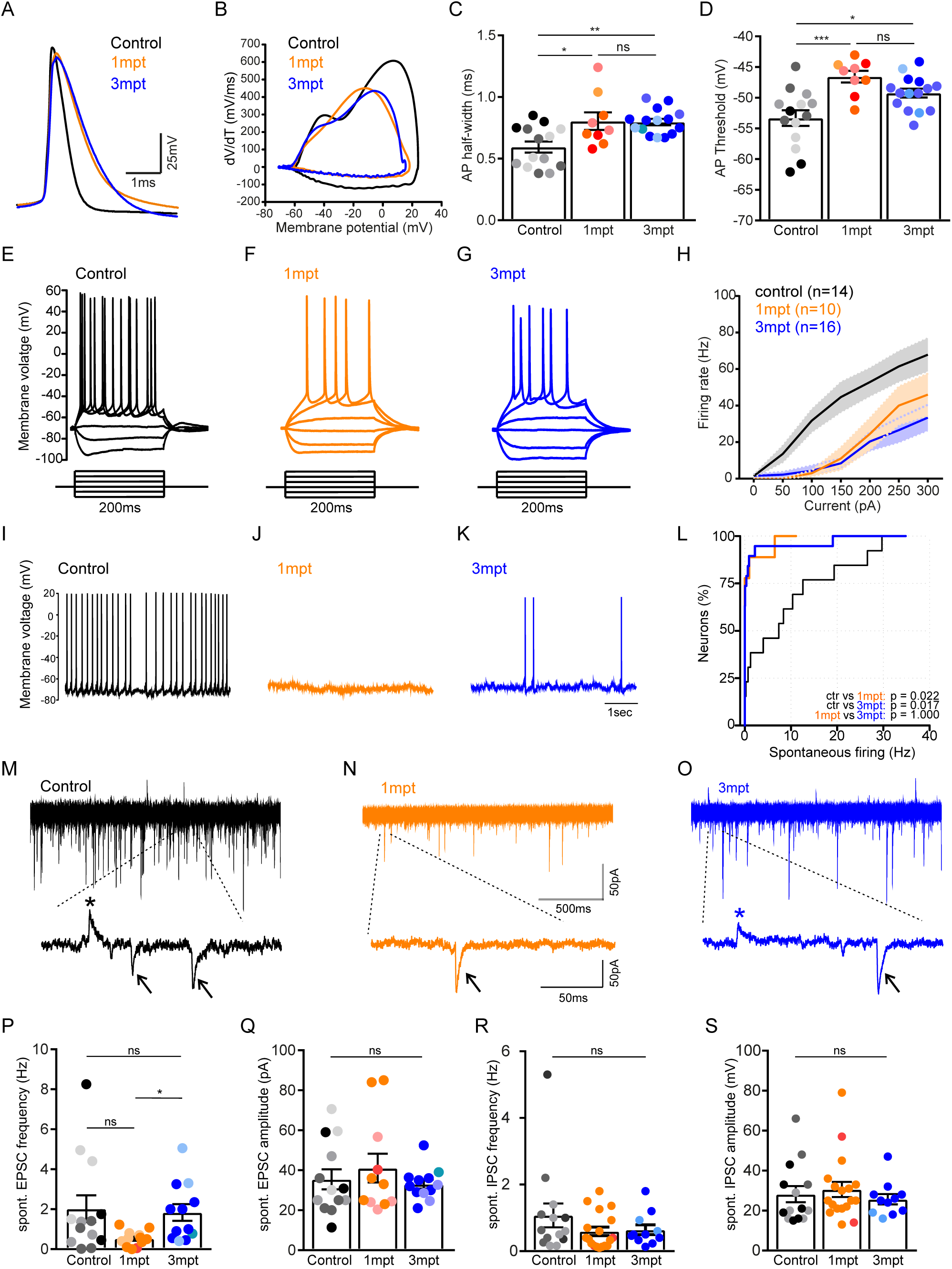
Action potentials and synaptic currents in tNs (A-D) Patch-clamp recording of action potentials (APs) of tN and control neurons in acute slices of injured brains. **(A)** Typical examples of AP in control (black), 1mpt (orange) and 3mpt (blue) neurons. **(B)** Phase plane plots of APs represent the rate of change of membrane potential (dV/dt) against the membrane potential (V) itself, illustrating the dynamics of the AP’s phases like depolarization and repolarization. **(C, D)** Quantifications of AP half-width and AP voltage thresholds (mice = 3-4, neurons (n) = 9-15). **(E-G)** Voltage traces in response to current injection (50pA steps) in control **(E)**, 1mpt **(F)**, and 3mpt **(G)** neurons. **(H)** Quantification of input-output functions in response to current injection. **(I-K)** Raw traces of spontaneous firing of action potentials in control **(I)**, 1mpt **(J)**, 3mpt **(K)**, and cumulative distributions **(L)** (mice = 3-4, neurons (n) = 10-16). **(M-S)** Whole-cell recording of spontaneous synaptic currents in tN and control neurons in acute brain slices. Raw traces of control **(M)**, 1mpt **(N)** and 3mpt **(O)** neurons. Insets show sIPSCs (asterisks) and sEPSCs (arrows). **(P-Q)** Quantification of sEPSC frequency **(P)**, and amplitude **(Q)** (mice = 3-4, neurons (n) = 11-13). **(R-S)** Quantification of sIPSC frequency **(R)**, and amplitude **(S)** performed at membrane voltage depolarized (-50mV) compared to resting membrane voltages (mice = 2-3, neurons (n) = 11-18). All data are presented as means ± SEM. Data obtained from different mice is denoted in different shades. ns p >0.05, * p <0.05, ** p <0.005, *** p <0.0005 by one-way ANOVA with Bonferroni test (C, D), Pearson Chi-squared test (L), and Kruskal-Wallis test with Dunn’s test (P-S).

As tNs were largely silent, we next investigated if this is due to a lack of functional synaptic input, by recording spontaneous excitatory and inhibitory postsynaptic currents (sEPSC and sIPSC) (Fig. 3M-O). We observed that sEPSC frequencies in tNs were reduced compared to neighboring endogenous neurons at 1mpt, but recovered to levels similar to endogenous neurons at 3mpt (Fig. 3P). This pattern corroborated our observations at the ultrastructural level with many spines being empty initially which decreased at 3mpt (Fig. 2B, C). Thus, these independent techniques both demonstrate that some aspects of tN synaptic integration are improved by 3mpt. Current amplitudes of sEPSCs were measured as a proxy for postsynaptic receptor quantity and quality, and were similar between tNs at 1mpt and also at 3mpt compared to endogenous neurons (Fig. 3Q). sIPSCs occurred as upward deflections in the current trace due to negatively charged chloride ions entering the cell. The sIPSCs show the typical slower decay time constants compared to sEPSCs, but their amplitudes were quite small (Fig. 3M-O insets) due to the weak driving force for inhibition at the neurons’ resting potential. To quantify sIPSCs, neurons were clamped to more depolarized values of −50 mV. Overall, sIPSCs occurred less frequently than sEPSC, but no significant changes in frequency or amplitude were observed across the three conditions (Fig. 3R, S). Thus, the many symmetrical synapses observed on the tNs in ultrastructural analysis (Fig. 2D) may not all be functional. This observation further hints at the immaturity of tNs, where during development many symmetrical synapses lack inhibitory functionality {*28*}, and some even proceed to mature into asymmetrical excitatory synapses {*29*}. During these recordings, we also used afferent synaptic stimulation to elicit short-latency evoked EPSCs (eEPSCs). Both, monosynaptic (Fig. S3E) and disynaptic (Fig. S3F) excitatory inputs could be evoked in neurons from all three groups, suggesting that generally tNs are equally well integrated into local microcircuits as the endogenous neurons. There is also an increased number of tNs at 3mpt exhibiting evoked EPSCs in (compared to 1mpt; Fig. S3G), similar to the pattern observed for spine maturation (Fig. 1E), ultrastructural analysis (Fig. 2B, C), and sEPSCs of the tNs (Fig. 3P). Taken together, these results show that many functional aspects of synaptic connectivity mature between 1 and 3mpt, even though many tNs remain silent.

In summary, our multimodal analysis revealed a consistent pattern of stunted maturation in the SWI environment: tNs maintain excess spines (Fig. 1C), many of which lack presynaptic partners (Fig. 2B); their synapses are disproportionately located on dendritic shafts rather than spines (Fig. 2C); and functionally, the majority of tNs remain silent (Fig. 3J-L). While partial maturation occurs between 1 and 3mpt across all readouts, tNs remain measurably distinct from endogenous neurons. This convergent evidence raised the question of what aspect of the host environment might be driving this persistent immaturity.

Spatial transcriptomics reveal activated microglia signature at the transplantation site To better extend the interaction between tNs and the surrounding cell types other than neurons they are connected with, we used spatial transcriptomics to investigate the molecular and cellular hallmarks of the environment exposed to SWI and transplantation in an unbiased manner. Towards this aim, we used spatial transcriptomics. Gene expression in sections of the cerebral cortex at 1mpt {*30*} was compared to sections with only a SWI without transplant (1mpi) (Fig. 4A). We integrated both datasets using Harmony {*31*} to correct for any batch effects and identified 18 clusters (Fig. 4B). Notably, two of the three clusters differing between the conditions corresponded to different anatomical structures present due to slight differences in the section level of the sample (Fig. S4A), while cluster 16 was enriched at the transplantation site (Fig. 4B, Fig. S5B). Since Visium spatial transcriptomics is spot-based, i.e. capturing transcripts in a 50µm diameter, each spot can contain transcripts from different cell types. To gain more insight into the cellular components of cluster 16, we performed a cellular component analysis using Cell Marker 2024 {*32*} on the significantly upregulated transcripts detected in cluster 16. The only significant cell identities distinguished were of immune cell origin, with microglia as the major cell type in the cluster (Fig. 4C), suggesting that this cluster is due to the altered accumulation and/or expression hallmarks of microglia. We then performed a gene ontology (GO) analysis and observed immune responses as significant GO terms upregulated in cluster 16 (Fig. 4D).

**Figure 4.**
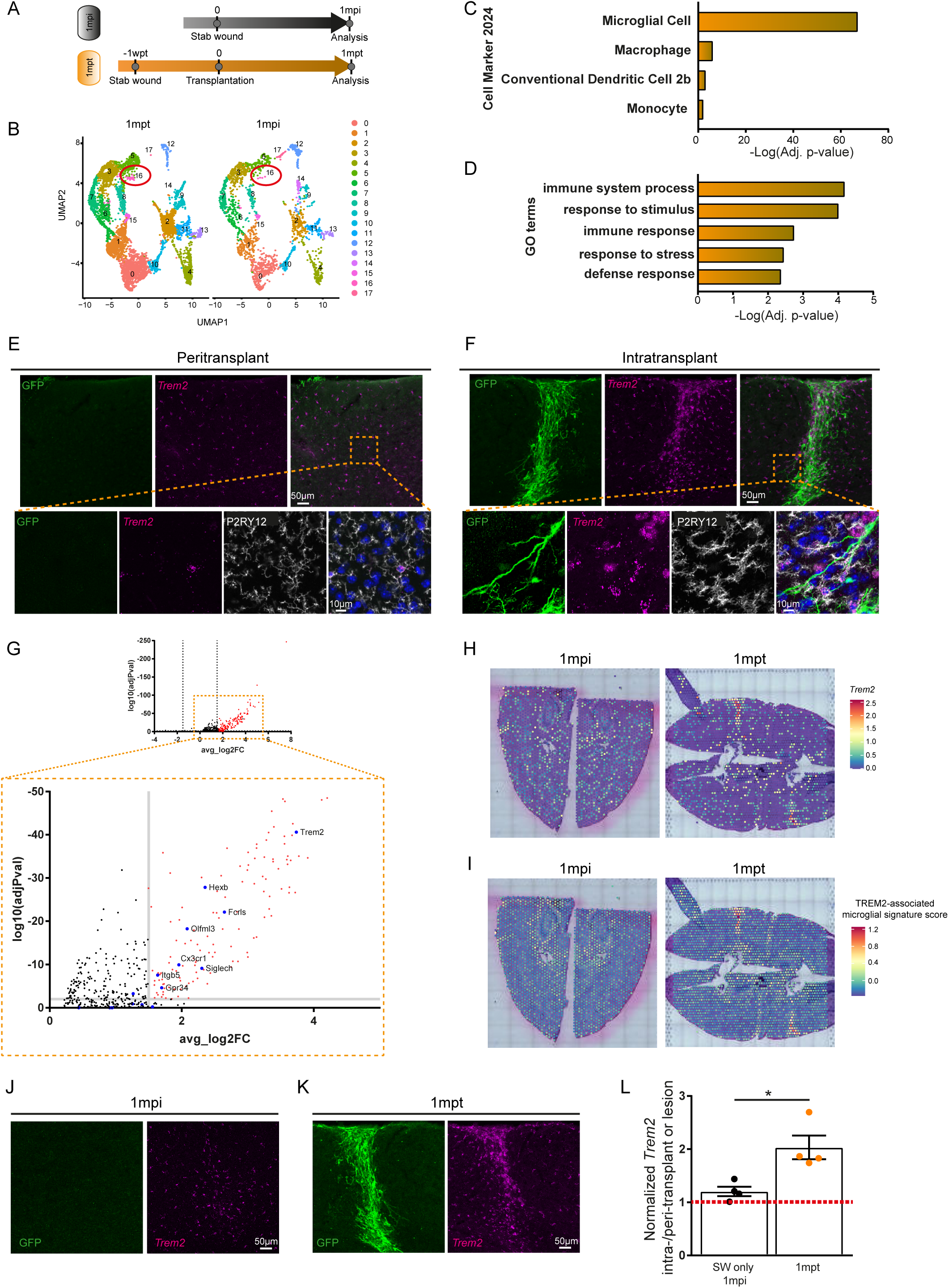
lncreased levels of *Treml* at the transplantation site. **(A)** Experimental setup used for spatial transcriptomics. **(B)** Uniform Manifold Approximation and Projection (UMAP) representation of Visium spatial transcriptomic datasets. Each point corresponds to a spatially resolved spot, colored by clusters. Cluster 16 is highly enriched in the 1mpt (stab wound followed by transplantation) dataset versus 1mpi (stab wound only). **(C)** Significant cell types identified in cluster 16 using Cell Marker 2024 (*32*). **(D)** Significant GO terms (*89*) identified upregulated in cluster 16. **(E-F)** Images showing the expression of *Trem2* (magenta) at the peritransplant (E) and intratransplant (F) regions. Transplanted cells express GFP (green). Presence of microglia was assessed using the P2RY12 marker (white). **(G)** Volcano plot of all the transcripts in cluster 16 (top). Inset (bottom) illustrates the significant microglia sensome transcripts detected in cluster 16. **(H)** Spatial transcript plot illustrating *Trem2* expression. **(I)** Spatial transcript plot illustrating the module score of TREM2-associated microglial signature. **(J-K)** Images showing the expression of *Trem2* (magenta) at the injury only (J; 1mpi), or injury + transplantation (K; 1mpt) or site. Transplanted cells express GFP (green). **(L)** Graph comparing the normalized mean gray values of *Trem2* expression at 1mpt, and 1mpi. (Mann-Whitney test, * p<0.05; mice (n) = 4). All data are presented as means ± SEM.

### *Trem2* remains activated at transplantation site

To determine, if and how microglia may enrich at the transplant site and how they interact with the tNs, we performed 2-photon live imaging experiments using *Cx3cr1^GFP^* mice to visualize microglia and labelled the tNs with a red fluorophore (FP). When imaging the transplantation site 1 week after transplantation (wpt), some microglia had a rather amoeboid morphology, indicating their high reactivity (Fig. S4C-E). However, this morphology changed by 2wpt and by 3wpt microglial sphericity was reduced and stabilized (Fig. S4D-E). Interestingly, a disparity in microglia morphology at 1mpt and 3mpt could be observed depending on the imaged region (Fig. S4F, G). Around the transplant, microglia were more ramified, i.e. less activated, while within the transplant they had a more reactive-like morphology (Fig. S4G). This observation could relate to the changes seen in microglia-related gene expression in the transplanted SWI condition and may have an influence on synaptic connectivity. Indeed, the differential morphology of microglia could be corroborated in brain sections (Fig. 4E; F, Fig. S4H), where microglia at the transplant core were identified as having shorter processes (Fig. S4I), and in general a less ramified morphology (Fig. S4J-L), suggesting persistent microglial activation in the transplant region.

Given the increasing evidence for persistent microglia activation at the transplant site, we returned to the spatial transcriptome data exploring the genes up-regulated in cluster 16 with a focus on the microglia sensome, a set of ligands and receptors on their surface, defining the capacity of microglia to sense the environment {*33*}. Amongst the top 25 microglia-specific sensome genes, we found 8 (*Trem2*, *Hexb*, *Fcrls*, *Olfml3*, *Cx3cr1*, *Siglech*, *Itgb5*, *Gpr34*) to be significantly upregulated in cluster 16, with *Trem2* as the highest up-regulated and most significant sensome gene (Fig. 4G). Additionally, *Trem2* was highly enriched at the transplantation site in the spatial plots (Fig. 4H). To validate this observation, we performed RNAscope of *Trem2* on sections of the cerebral cortex (Fig. 4E, F, J, K). This showed that *Trem2* was significantly upregulated in the transplant core compared to stab wound only at 1mpi/mpt (Fig. 4J-L). Moreover, *Trem2* shows an approximately 2-fold higher expression in the intratransplant compared to the peritransplant region at 1 and 3mpt (Fig. S4M-N).

We then assessed if TREM2-associated microglia signature is persisting at the transplantation site using spatial transcriptomics. This signature was shown to be lacking in *Trem2^-/-^* mice {*34*}. Thus, we created a module score with the top 10 genes (*Trem2*, *Cst7*, *Spp1*, *Lpl*, *Gpnmb*, *Ank*, *Itgax*, *Ctsl*, *Ctsz*, *Cd68*, *Axl*) of this TREM2-associated microglia signature, and found it to be highly elevated in the transplant sections compared to 1mpi sections (Fig. S5A), particularly in cluster 16 (Fig. S5B). Indeed, plotting this module score spatially exhibited localized elevated levels at the transplantation site (Fig. 4I).

These data indicate that microglia around the transplantation site are chronically activated as detected by their morphology and gene expression with a prominent TREM2-mediated signature, suggesting that the presence of tNs in the SWI aggravates a TREM2-associated inflammatory environment.

### TREM2 regulates tN circuit remodeling

Microglia are known circuit remodelers {*11–13*, *17*}. Additionally, prolonged activation of TREM2 in microglia has been associated with a dysregulation of this microglia ability {*34*, *35*}. Thus, we checked if microglia were altered at the transplantation site in the cortex of *Trem2^-/-^* mice. Morphological analysis revealed that microglia at the transplantation site in *Trem2^-/-^* cortex exhibited fewer morphological changes compared to those in wildtype (wt) hosts (Fig. S4H-L), indicating that transplanting into a TREM2-free environment changes microglial reactivity.

Having established that microglia at the transplant site show reduced reactivity in *Trem2⁻^/^⁻* cortex, we next asked whether this altered microglial state affects the brain-wide input connectivity of tNs. To map which neurons in the brain form synaptic connections onto tNs, we used monosynaptic rabies virus (RABV) tracing, a well-established technique that labels presynaptic neurons providing direct synaptic input to a defined starter cell population {*2*, *7*, *8*}. Prior to transplantation, e14 cortex cells were transduced *in vitro* with a retrovirus encoding for TVA, the receptor recognized by an EnvA-pseudotyped RABV, ensuring that RABV selectively infects tNs and spreads retrogradely to their direct presynaptic partners, but no further. The retrovirus encoding TVA and the glycoprotein also includes a sequence for a fluorescent protein (XFP), to identify the transduced tNs. Cells were transplanted 1 week after SWI as in all other experiments and 1 week prior to the 1mpt and 3mpt timepoints (Fig. 5A), the EnvA-pseudotyped SAD-ΔG-XFP RABV (with a fluorophore different from the one of the tNs) was injected into the transplant site. This allowed the identification of tNs by the expression of the retrovirus XFP, their presynaptic partners by the expression of the RABV XFP, and the starter cell population by the expression of both XFPs (Fig. 5B).

**Figure 5.**
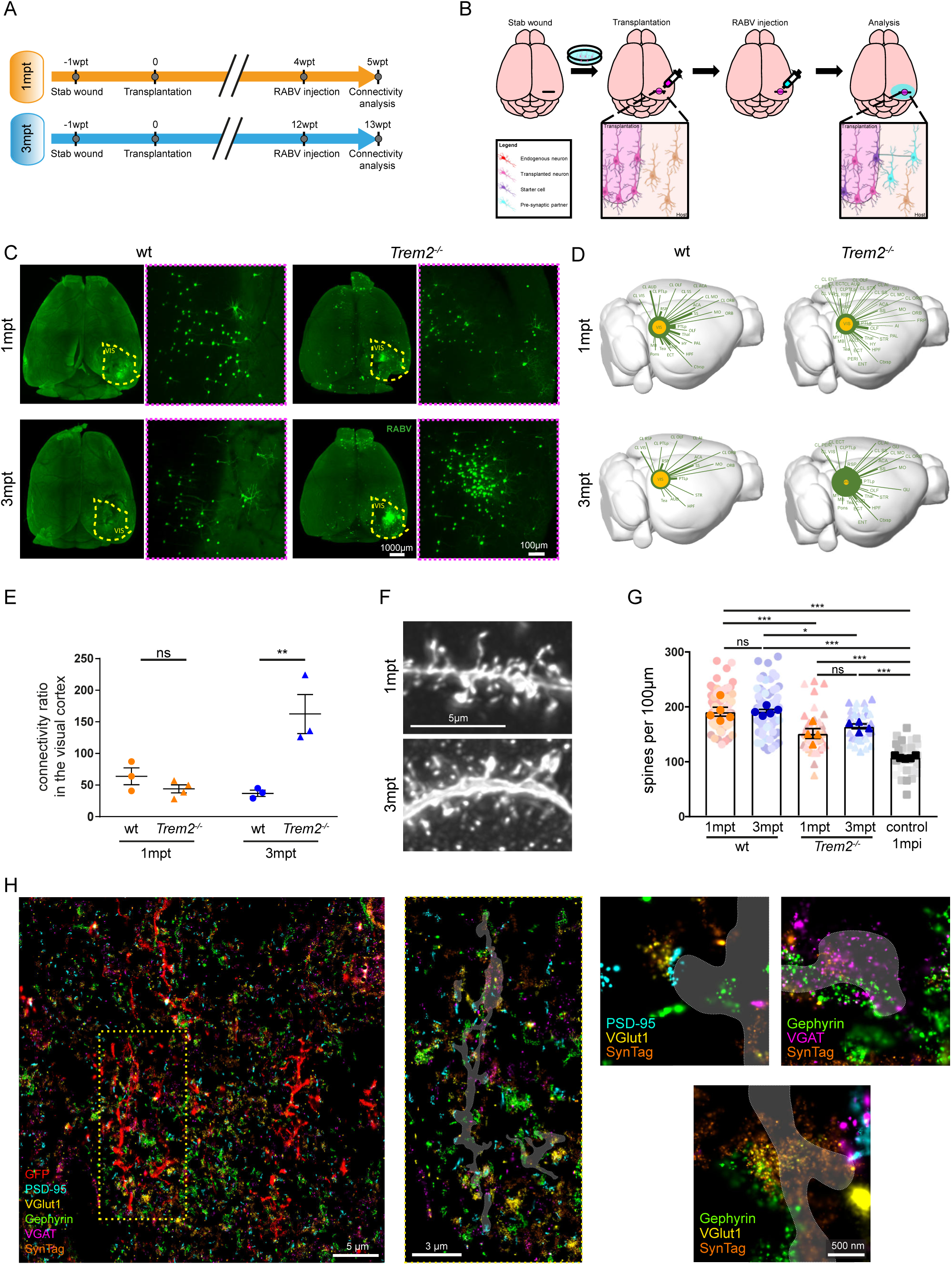
TREM2 is a circuit remodeler for tNs. **(A)** RABV-tracing of tNs experimental setup and **(B)** the quantitative circuit mapping using monosynaptic retrograde RABV tracing. **(C)** Whole brain images obtained using light sheet microscopy showing tN presynaptic partners, traced using monosynaptic retrograde RABV. Insets (dotted magenta boxes) contain a zoom-in on the light sheet images at the visual cortex. **(D)** 3D connectograms with line thickness depicting the connectivity ratio for a given brain region of Rabies-traced brain. wt (left) and *Trem2^-/-^* (right) animals at 1mpt (top) and 3mpt (bottom). **(E)** Quantification of connectivity ratios (input neurons/starter neurons) in the visual cortex of wt and *Trem2^-/-^* mice at 1mpt and 3mpt (mice = 3-4). **(F-G)** Images of dendritic processes with spines from tNs in *Trem2^-/-^* at 1mpt (top) and 3mpt (bottom) **(F)**, and spine density quantifications **(G)**. Data from wt and control brains were taken from Figure 1F (mice (n) = 4 - 5, dendrites = 31-38). **(H)** six-plex DNA-PAINT imaging region, depicting the GFP-positive tN (red), PSD95 (cyan), vGLUT1 (yellow), gephyrin (green), vGAT (magenta), SynTag (orange). Inset of a tN dendrite from (H) and zoom-ins depicting excitatory (top), inhibitory (middle), and mixed (bottom) synapseIn graphs, solid colors represent per-animal means; lighter shades represent individual measurements, with distinct shades denoting each mouse. All data are presented as means ± SEM. All statistics were performed on the means of individual mice - ns p >0.05, * p <0.05, ** p <0.005, *** p <0.0005 by two-way ANOVA (E), and One-way ANOVA with Tukey’s multiple comparison test (G) performed on animals.

The brain-wide connectome of tNs was imaged by using whole brain clearing followed by light sheet microscopy (Fig. 5C) and presynaptic partners were then semi-automatically quantified. For each brain region, we calculated tN connectivity ratios by normalizing the presynaptic numbers in the brain region of interest to the number of starter cells (Fig. 5D, E, Fig. S5C-E, G). The most striking finding was a profound increase in the inputs from the ipsilateral visual cortex in *Trem2^-/-^* mice at 3mpt (Vis; Fig. 5C-E, Fig. S5G), while this local input was reduced between 1 and 3mpt in wt (Fig. 5E; see also (*7*)). Comparing tN connectivity ratios at 1mpt and 3mpt across brain regions between wt and *Trem2^-/-^* mice, the visual cortex (Vis) was the only region showing significant increase (Fig. S5C-E). Conversely, inputs from several distant brain regions were lost at 3mpt (Fig. 5D). However, the remaining connected brain regions in *Trem2^-/-^* mice more closely matched the regions normally connected to the visual cortex {*2*}, suggesting improved circuit specificity. Thus, the lack of TREM2 allows local inputs to further increase between 1-3mpt, while they are rather reduced in the wt between 1 and 3mpt. Importantly, no significant difference in connectivity was observed between *Trem2^-/-^* and wt at 1mpt (Fig. 5E), indicating that TREM2 does not affect the initial establishment of connections but rather their subsequent refinement between 1 and 3mpt. To ensure that the connectivity results were not biased by the number of starter cells, we performed a Pearson correlation test between the number of starter cells and the number of presynaptic partners (Fig. S5F). As seen before {*7*, *8*}, also in this data set there is no significant correlation between the number of starter cells and the input connectome (Fig. S5F). Altogether, these results identify TREM2 as a mediator of tN circuit remodeling, specifically limiting the consolidation of local synaptic inputs during the refinement phase between 1 and 3mpt.

### Spine density of tNs in *Trem2^-/-^* animals is partially restored

Given that tNs in wt SWI exhibited a persistent 2-fold increase in spine density (Fig. 1C), we next asked whether the absence of TREM2 would normalize this. Building on the connectivity analysis results, we proceeded to analyze spine density of tNs in *Trem2^-/-^* mice, based on GFP immunostaining (Fig. 5F). Similar to our observations in the wt environment (Fig. 1C; Fig. 5G), no significant changes in spine density were observed between 1mpt (152 ± 39 spines/100µm) and 3mpt (164 ± 29 spines/100µm; Fig.5G), in *Trem2^-/-^* mice. Although the spine density of tNs in *Trem2^-/-^* mice was still significantly increased when compared to the control endogenous neurons in the intact brain, these tNs exhibited a lower spine density than the tNs in the wt condition (Fig. 5G). Quantitative analysis of the different spine types of tNs (Fig. S5H-M) showed no significant difference between 1mpt and 3mpt in the *Trem2^-/-^* mice, reaching similar levels of mushroom and filopodia spines as in tNs at 3mpt in wt brains (Fig. 1E, F). Altogether, these results suggest that the synaptic maturation of tNs is accelerated and partially normalized toward control levels in the *Trem2^-/-^* environment.

### Increase in complete chemical synapses of tNs in *Trem2^-/-^* environment

To explore the synaptic integration of tNs in the *Trem2^-/-^* environment beyond morphological analysis, we probed their neurochemical composition using DNA-PAINT analysis (Fig. 5H). We observed that of the total synapses analyzed (n=164), there was an increase in full synapses. In fact, both full excitatory (53.1%) and inhibitory (33.3%) chemical synapses almost doubled in percentage. However, a large percentage of synapses, both inhibitory and excitatory, were still empty. The many empty spines we observe may reflect that we are only probing specific proteins, as a presynapse could still be present, but with a different molecular machinery. Altogether these results further suggest that tNs in the *Trem2^-/-^* environment become more adequately connected compared to the wt.

### Functional electrophysiological properties of tNs are restored at 3mpt in *Trem2*^-/-^

Given that tNs in wt SWI remained largely silent with immature action potential properties (Fig. 3), we next assessed the functionality of tNs and their integration in the *Trem2^-/-^* environment. We performed patch-clamp recordings from acute slices of cerebral cortices, as in the wt setting (Fig. 3). We observed that tNs in the *Trem2^-/-^* environment also reliably generated action potentials with long half-width and more depolarized thresholds. However, contrary to tNs in the wt, action potential half-width thresholds were restored to control values from neighboring endogenous neurons at 3mpt (Fig. 6A-D). Furthermore, in response to increasing depolarizing current injections, all three conditions generated increasing number of action potentials (Fig. 6E), with tN firing rates similar to those of endogenous neurons at the SWI site at both 1mpt and 3mpt (Fig. 6E, F). This restoration of functional properties was also observed when we measured spontaneous action potentials (Fig. 6G), where no significant difference was observed between the endogenous neurons and tNs (Fig. 6H). In summary, tNs in the *Trem2^-/-^* environment reached an intrinsic functional maturation comparable to the neighboring endogenous neurons. This is very different from tNs in a wt environment (Fig. 3J, K), showing the potent influence of the environment mediated by TREM2 on functional maturation of tNs.

**Figure 6.**
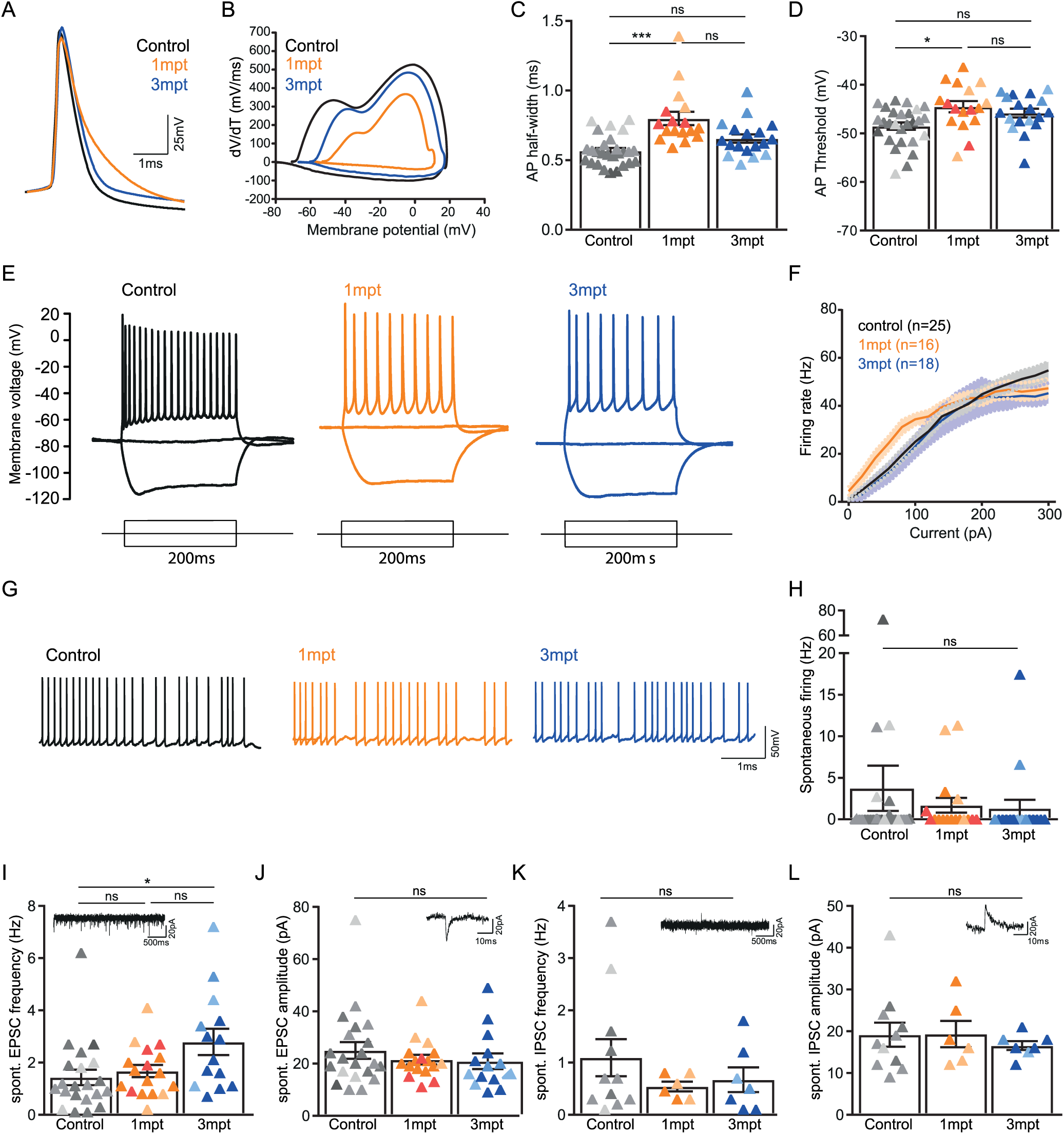
Functional electrophysiological properties of tNs are rescued in *Treml^-/-^* (A-D) Patch-clamp recording of action potentials (APs) of tN and control neurons in acute slices of injured brains. **(A)** APs in control (black), 1mpt (orange) and 3mpt (blue) neurons. **(B)** Phase plane represent the rate of change of membrane potential (dV/dt) against the membrane potential (V) itself, illustrating the dynamics of the AP’s phases like depolarization and repolarization. **(C, D)** Quantifications of AP half-width and AP voltage thresholds (mice = 3-4, neurons (n) = 17-27). **(E)** Voltage traces in response to current injection (50pA steps) in control (black), 1mpt (orange), and 3mpt (blue) neurons. **(F)** Quantification of input-output functions in response to current injection. **(G)** Raw traces of spontaneous firing of action potentials in control (black), 1mpt (orange), 3mpt (blue), and their plotted values **(H)** (mice = 3-4, neurons (n) = 17-27). **(l-J)** Quantifiction of sEPSC frequency **(l)**, and amplitude **(J)** (mice = 3-4, neurons (n)= 17-27). **(K-L)** Quantification of sIPSC frequency **(K)**, and amplitude **(L)** performed at membrane voltages depolarized (-50mV) compared to resting membrane voltages (mice = 2-3, neurons (n) = 6-11). Insets show example traces at two different time scales. All data are presented as means ± SEM. Data obtained from different mice is denoted in different shades. ns p >0.05, * p <0.05, ** p <0.005, *** p <0.0005 Kruskal-Wallis test with Dunn’s test.

Next, we measured the synaptic currents of tNs in *Trem2^-/-^*. Already at 1mpt similar sEPSC frequencies were recorded in tNs compared to control neurons. The sEPSC frequency increased even further (Fig. 6I), matching the RABV connectivity analysis (Fig. 5E). The sEPSC amplitudes did not differ between the different groups (Fig. 6J). Conversely, sIPSCs did not show any difference between tNs and endogenous adjacent neurons in frequency (Fig. 6K) or amplitude (Fig. 6L). Overall, these results confirm that unlike in the wt environment, the tNs in the *Trem2^-/-^* environment have improved maturation and integration trajectory with functional parameters restored to endogenous levels.

### Spatial transcriptome analysis in *Trem2^-/-^* cortex reveals a more permissive environment for tN synaptic integration

To understand the molecular underpinnings of the synaptic changes observed above in an unbiased manner, we performed a 10x Visium HD spatial transcriptome analysis of wt and *Trem2^−/−^*animals at 1 and 3 mpt. We applied bin2cell to bring the bin-based Visium HD data to single cell resolution and isolated GFP-positive cells (Fig. 7A; Fig. S6A, B). Unsupervised clustering on the combined four datasets revealed that clusters were mostly corresponding to brain region and cell type and not driven by batch effects (Fig. 7A, Fig. S6A). Nevertheless, the transplanted area in each condition formed a distinct cluster (Fig. S6A), indicating a clear transcriptional signature distinguishing the transplant site from surrounding endogenous tissue. Differential gene expression (DEG) analysis revealed a major shift in glia-associated transcripts at the transplantation site in wt compared with *Trem2^−/−^* brains (Fig. 7B, Fig. S6C). Specifically, processes associated with glia predominated in the wt condition, whereas in *Trem2^−/−^* brains the most enriched processes related to the regulation of synaptic transmission and signaling (Fig. 7B, Fig. S6C). As expected, the TREM2-associated microglial signature was upregulated in wt but not in *Trem2^−/−^*brains (Fig. 7C).

**Figure 7.**
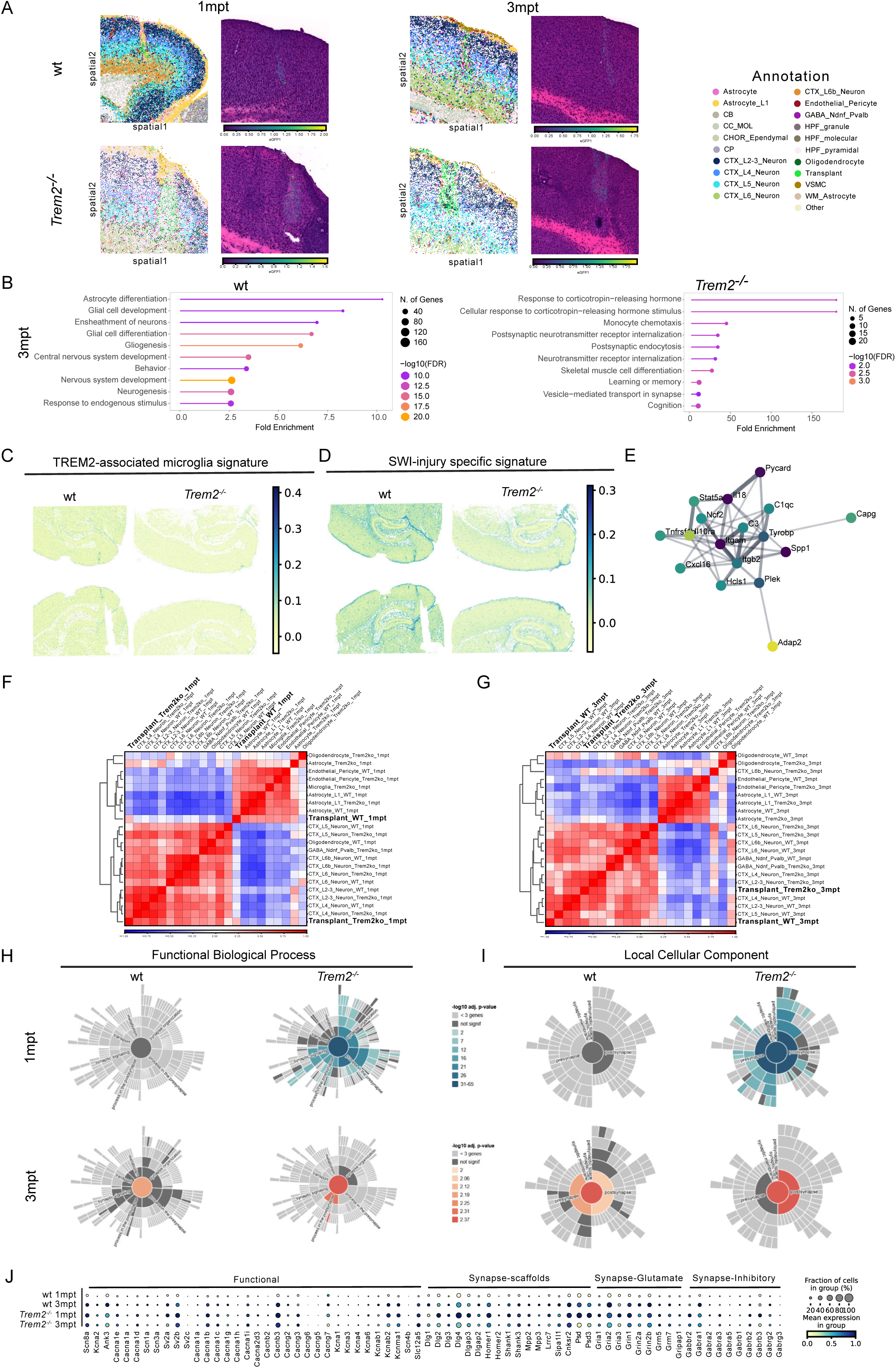
Spatial transcriptome analysis reveals reduced inflammatory signature and increased tN synaptic maturation in a *Treml^-/-^* environment. **(A)** Spatial plots overlaying cluster annotations (left) and presence of GFP signal (right), for the different conditions. **(B)** Significantly top 10 upregulated biological processes based on the DEGs in the transplant cluster in wt and *Trem2^-/-^* at 3mpt. **(C-E)** Different signature expressions in wt and *Trem2^-/-^* at the transplantation site. **(C)** TREM2-associated microglial signature, and **(D)** SWI-specific signature. **(E)** STRING analysis of transcripts elevated in the wt condition show DAP12 microglial causal network. Highlighted transcripts detected in our spatial dataset involved in the DAP12 microglial causal network. Network from wikipathways. **(F-G)** Correlation matrices of the annotated cell types in wt and *Trem2^-/-^* at 1mpt **(F)**, and 3mpt **(G)**. **(H-J)** Synaptic processes upregulated in tNs in wt and *Trem2^-/-^* brain at 1 and 3mpt of **(H)** functional biological processes, and **(I)** local cellular component. **(J)** Dot plot of transcripts associated with synapse functionality, scaffolds, excitatory and inhibitory synapses expressed by tNs of the different conditions.

To further characterize the injury environment surrounding tNs, we examined the SWI-specific proteomic profile we recently reported (*7*), which includes several inflammation-associated proteins. This profile was upregulated in wt brains but not in *Trem2^−/−^* brains (Fig. 7D), partly owing to a reduction in complement factors in the *Trem2^−/−^* environment. Moreover, one of the most significantly upregulated pathways in wt brains was the DAP12 (Tyrobp) causal network (Fig. 7E, Fig. S6D), a key regulator of microglia activation [*36*). Transcripts both upstream and downstream of DAP12 were upregulated in wt but not in *Trem2^−/−^* brains (Fig. S6E), indicating that microglia in *Trem2^−/−^* mice are not chronically primed toward activation.

Altogether, these data indicate that the *Trem2^−/−^* cortex harbors a reduced complement and neuroinflammatory signature compared with wt, providing evidence that modulation of the injury environment surrounding tNs can render it more permissive for the maturation and integration of tN synapses. We therefore proceeded to investigate the transcriptional profile of the tNs themselves.

### Spatial transcriptome analysis in *Trem2^-/-^* cortex reveals tNs are more synaptically mature than tNs in wt environment

Next, we investigated if this change in environment results in any differences in the tN transcriptome. Correlation analysis across all annotated cell types revealed that tNs in wt tissue clustered most closely with astrocytes at 1 mpt, whereas tNs in *Trem2^−/−^* brains already clustered with endogenous cortical neurons at the same time point (Fig. 7F). By 3 mpt, tNs in both genotypes clustered with endogenous neurons (Fig. 7G); however, tNs in wt hosts at 3 mpt remained transcriptionally closer to *Trem2^−/−^* tNs at 1 mpt (Fig. S6F), suggesting a delayed maturation trajectory in the wt environment.

To examine synaptic maturation specifically, we queried the GFP-positive cells against the SynGO database {*37*}. Many synapse-associated genes linked to synapse maturity were differentially upregulated in *Trem2^−/−^* but not in wt tNs (Fig. 7H, I). Notably, tNs in wt brains showed no significant upregulation of synaptic processes at 1 mpt and only minimal enrichment at 3 mpt (Fig. 7H, I), indicating that although synaptic maturation occurs between 1 and 3 mpt in the wt condition, it does not reach the levels observed in the TREM2-deficient condition. Consistent with this, transcripts encoding synaptic scaffolding proteins, both excitatory and inhibitory, and functional synaptic transcripts were generally increased in wt tNs between 1 and 3 mpt (Fig. 7J), confirming a slow but ongoing maturation process.

Altogether, these data suggest that tNs in the wt environment undergo gradual synaptic maturation between 1 and 3 mpt but remain less mature than those in *Trem2^−/−^* brains. Furthermore, by 3mpt in the *Trem2^-/-^* environment, the transcriptional identity of tNs is highly correlated to that of endogenous upper layer cortical neurons, consistent with our earlier observations at the functional and connectivity levels.

## Discussion

Using a multimodal approach spanning morphological, ultrastructural, and functional analyses, we demonstrated that tNs in the SWI environment exhibit a consistent pattern of stunted synaptic maturation across all readouts. Spatial transcriptomics identified a persistent TREM2-associated microglia signature at the transplant site as a key environmental factor, and transplantation into *Trem2⁻^/^⁻* mice largely rescued these deficits. These findings establish that the inflammatory host environment, not intrinsic neuronal limitations, is the primary bottleneck for tN integration; thus, highlighting the crucial importance of the injury-mediated environment for tN maturation and integration. These data demonstrate the need to not only focus on optimizing protocols for neuronal differentiation prior to transplantation, but also including strategies to regulate the environment surrounding them.

### Integration of new neurons in the adult brain

We used the SWI model to examine how synapses of tNs mature and integrate in an inflamed adult brain using a deep multimodal synapse analysis. This analysis enabled us to capture a holistic picture of the tN synaptic milieu and unlocked insights that would have otherwise been missed. Much to our surprise, we have observed that when transplanting in the SWI in the wt condition, the different synapse read-outs, i.e. morphological vs. chemical vs. functional, do not necessarily align with each other. For example, we find a large proportion of synapses on shafts and with symmetric pre- and postsynapses, yet no clear increase in inhibitory synaptic potentials (Table 1). However, most of the differences we found in tNs’ synaptic integration compared to endogenous neurons, such as higher number of spines, more empty spines without a presynapse, more synapses on shafts, larger half-width of APs and less spontaneous firing activity point to a certain degree of immaturity in functional and synaptic maturation. Notably, synaptic potentials of tNs are similar to endogenous neurons in the same injury environment in regard to their amplitude and frequency (Table 1), but their APs are still immature with a larger half-width and lower frequency when elicited. This is reminiscent to the “first listen, then fire” mode of neuronal integration in adult olfactory bulb (OB) neurogenesis. OB interneurons born in development fire APs around the time they finish migrating {*38*}, while adult-born OB interneurons acquire sodium conductance and fire APs at later maturational stages, after receiving synaptic input on their somata and later at their proximal dendrites {*38–40*}. However, functional maturation of adult generated OB interneurons is achieved within about 1 month, while tNs in SWI still show many of these immature hallmarks, albeit in reduced amount (Table 1], at 3mpt. This may have dire consequence for their function in the network and achieving further aspects of functional maturation, such as axonal myelination {*41*}.

**Table 1.** Synaptic multi-modal readouts of transplanted neurons in the Stab wound injury wildtype mouse brains.

| **Synaptic readout** | **Overall phenotype** | **Additional Information** | **From 1-3mpt** | **conclusion** |
| --- | --- | --- | --- | --- |
| Spines | Increased in number | More filopodia spines, decrease between 1-3mpt, but still higher than eNs | Increased mushroom spines at 3months, same level as endogenous neurons | Immature state |
| EM | Empty spines (ES);<br>Spines on shafts (SoS) |  | ES decrease, but remain more; SoS decrease, but remain more | Immature state |
| Neurochemical staining | Empty postsynapses |  | Increase of excitatory synapses | Partial immature state |
| Electrophysiology: synaptic potentials | Normal frequency & amplitude at 3mpt | Amplitude of IPSCs&EPSCs normal already at 1mpt | Spontaneous EPSC frequency increases | Mature, But fewer cells with sEPSCs |
| Electrophysiology: action potentials | Larger halfwidth; fewer evoked and much fewer spontaneous APs |  | A few more spontaneous APs | Persistent immature state |
| RABV tracing | Innervation from adequate regions |  | Pruning and loss of local input neurons |  |

### Immature synaptic integration and physiology of tNs in the injured brain

Generally, neuronal maturation proceeds well after transplantation into SWI condition acquiring pyramidal neuron hallmarks and molecular markers of cortical neurons {*2–4*, *7*}. However, by examining other synaptic parameters, we observe some aberrations, such as continued exuberant spine density or a high level of filopodia spines. Increased spine density is reminiscent of developing neurons, where filopodia are the main spine type {*21*}. Filopodia are long, highly motile dendritic protrusions designed to search for presynaptic partners and consolidate them into stable synapses {*21*}. Of note, tNs in wt SWI exhibited an immature spine type profile, which partially resolved at 3mpt. Yet, filopodia percentages remained 2-fold higher compared to endogenous neurons. Additionally, in the adult brain, spine synapses make up around 85% of the synapses {*9*}. In contrast, shaft synapses predominate the synaptic landscape in early development {*21*, *42*}. Using 3D-EM, our data show that almost 70% of the synapses on tNs at 1mpt are in fact shaft synapses, indicating their immaturity. This improves at 3mpt with 42% synapses on shafts of tNs, but remains higher compared to endogenous neurons (18%). In physiological development of the rodent cortex, neurons remain in this phase for the first two postnatal weeks {*21*}, however, it seems that tNs in the SWI are in this state for much prolonged time (3 months) or possibly perpetually. Another interesting aspect is that virtually all spines [>95%) in adult cortical neurons possess a presynapse {*43*}, while only half of the spines of tNs following SWI exhibit a presynapse, here referred to as “empty spine/synapse”. To date, the significance of the remaining non-synapse (empty) spines is not fully understood, but it was proposed that these spines in adulthood under physiological conditions could either be remnants of once synapse-containing spines or protrusions to increase the surface area for dendrites to form a synapse at a later timepoint {*43*}.

An important consideration is whether the abundant symmetric synapses observed on tNs by EM are truly inhibitory, or instead reflect immature excitatory synapses with thin PSDs that have not yet acquired the electron-dense, asymmetric morphology characteristic of mature glutamatergic synapses. Several lines of evidence support the latter interpretation. Our electrophysiology reveals no corresponding increase in sIPSC frequency and DNA-PAINT did not reveal excess amounts of inhibitory synapses. During early postnatal cortical development, a high proportion of synapses classified as symmetric by EM criteria show no immunoreactivity for inhibitory markers {*28*}, additionally it was reported that many morphologically symmetric synapses in developing tissue are in fact nascent excitatory synapses, which later develop into asymmetric synapses {*29*}. The convergence of our EM, DNA-PAINT, and electrophysiology data therefore favours the interpretation that the excess symmetric synapses on tNs reflect synaptic immaturity rather than a genuine shift toward inhibitory innervation, further supporting the conclusion that tN synaptic maturation is stunted in the SWI environment.

Given the importance of the host environment for tN integration {*7*}, we also compared the spine density of tNs in the SWI to the tNs in layer II/III neuron ablation model {*2*}, which is a targeted neuronal ablation model eliciting little to no inflammation {*44*}. In both these models, no major spine or dendritic pruning was observed between 1 and 3mpt. However, tNs in the SWI exhibited a 2-fold increase in dendritic spine density, when compared to tNs in the layer II/III neuron ablation model {*2*}. The spine density in the neuronal ablation model was similar to that observed in endogenous neurons in our study, indicating that the environment plays a crucial role in tN integration, and that a SWI environment lead to abnormal dendritic spine development of tNs. On a similar note, functional readouts using patch-clamp recordings of tNs in the SWI showed a degree of immature intrinsic properties, such as prolonged action potential-half width, reduced firing rate and the absence of spontaneous firing. Previous functional experiments of tNs in other mouse models, such as in the layer II/III neuron ablation model {*2*}, revealed that tNs exhibit prolonged sharpening of receptive field tuning, when compared to physiological neuronal development, however, by 3mpt, tNs had acquired receptive field properties identical to endogenous neurons {*2*}. The fact that different disease models, especially using immune-suppressed models, result in a different synaptic maturation of tNs is further corroborated by an EM study, where human induced pluripotent stem cell-derived tNs in a stroke model in nude rats were shown to have around 10% inhibitory and 90% excitatory synapses {*5*}. This is in concordance with others using EM analysis to assess the synapses of cortical pyramidal neurons in the postnatal and adult cortex across species, reporting synaptic ratios of 1:9 {*9*, *25*, *28*, *45*, *46*}. The 1:9 synaptic ratio is highlighted in {*25*}, where this synaptic ratio kept constant in the different species analyzed despite changes in neuronal composition. Particularly the inflammatory component in the SWI model is likely to result in increased inflammatory markers such as nitric oxide, which has been shown to suppress the KCC2 transporter, depolarize the chloride reversal potential and thereby turn truly hyperpolarizing inhibition into much smaller, merely shunting currents {*47*}. Thus, the excessive number of symmetric synapses observed may become active under certain physiological or stress conditions, which is why it is important to know that they are there. Notably, unlike in the SWI {*48*}, both the layer II/III neuronal ablation model {*44*} and the stroke model in nude rats {*49*}, where synapses had the normal 10:90 inhibitory to excitatory synapse composition {*5*}, lack inflammatory components. In fact, transplanting in *Trem2^-/-^* animals reverted many synaptic readouts to control levels, adding evidence that differences in functional and synaptic maturation of tNs in the wt condition is caused by inflammation.

### Role of microglia and TREM2 in tN maturation and integration

The convergent evidence of stunted maturation across all synaptic readouts, combined with the spatial transcriptomics discovery of persistent TREM2-associated microglia activation, pointed to microglia as the key mediators of the environmental influence on tN integration. Microglia have long been associated with non-immune activities, such as the sculpting of neuronal circuitry {*10–13*}, playing various roles in development and adulthood {*50*, *51*}. In fact, most aspects of synaptic integration and functional maturation are improved in the *Trem2^-/-^* environment (Table 2). A proteomic signature of inflammation aspects in the host tissue has been previously correlated with changes in tN connectivity {*7*, *8*}. Here we used spatial transcriptomics and unbiasedly discovered an increased microglia activation in the transplant condition, including a higher *Trem2* module. TREM2-signalling, when activated, pushes microglia to achieve a higher reactive stage {*34*}, which is thought to have detrimental effects, when activated chronically. Chronic inflammation stunts the development of new neurons, especially synapse maturation due to the cytokine milieu {*52*}. Additionally, modulating microglial reactivity results in altered neuronal development. For example, in *Cx3cr1^-/-^* animals, hippocampal neurons have a higher spine density and a delayed synapse maturation {*53*}, and during adult hippocampal neurogenesis, the presence of an inflammatory environment reduces the survival and recruitment of newborn granule cells {*54*} and skews their synaptic integration toward stronger inhibitory and weaker excitatory drives {*55*}. In development, reduction of microglia cells via migration to layer 1 and white matter has been shown to be crucial for adequate maturation of the differentiating neurons {*56*}, suggesting an adverse effect of microglia on these processes. However, also beneficial effects of microglia on neurogenesis have been reported that turn into adverse effects upon maternal immune activation {*57*, *58*}. Thus, the state of microglia influences neuronal maturation in development, but none of this has yet been linked to TREM2. Here we show a persistent microglia activation with a TREM2 signature in the transplant, and that TREM2 deletion normalizes all functional parameters of the tNs with normal AP firing frequency, AP half-width etc (Table 2), and improves the spine numbers and empty spines closer to endogenous neurons. In addition, the loss of TREM2 results in increased innervation as seen by a higher number of sEPSCs and local input neurons by RABV-tracing. While we cannot directly associate these effects to TREM2 on microglia, these findings reveal a novel role of TREM2 in limiting tN maturation, and a function in limiting tN synaptic integration well consistent with the classical role of TREM2 in circuit refinement in physiological {*10*, *59*} and pathological conditions {*14–16*, *60*}.

**Table 2.** Synaptic multi-modal readouts of transplanted neurons in the Stab wound injury *Trem2^-/-^* mouse brains.

| **Synaptic readout** | **Overall phenotype** | **Additional Information** | **From 1-3mpt** | **conclusion** |
| --- | --- | --- | --- | --- |
| Spines | Increased in number | Less increased compared to WT host | No change | Less immature state |
| Neurochemical staining | Less empty spines (ES); |  | Not analyzed | Less immature state |
| Electrophysiology: synaptic potentials | Normal frequency & amplitude at 1mpt | Amplitude of IPSCs&EPSCs normal already at 1mpt | Spontaneous EPSC frequency increases above control | Mature, More sEPSCs At 3mpt |
| Electrophysiology: action potentials | Larger halfwidth at 1mpt; normal evoked spontaneous AP frequency |  | Halfwidth normal at 3mpt | Mature |
| RABV tracing | Innervation from adequate regions at 1mpt |  | Excessive local innervation at 3mpt | Too little pruning |

As many studies probe tN integration by monosynaptic Rabies virus tracing, we also explored how this brain-wide input connectome would be shaped by the *Trem2^-/-^* environment. We observed an increase in connectivity levels of the local inputs between 1 and 3mpt in the *Trem2^-/-^* hosts, but not in a wt environment, where inputs were rather reduced (Fig. 5). Importantly, long-range inputs of tNs were to some extent decreased between 1 and 3mpt in the *Trem2^-/-^* hosts. Although, the lost long-range inputs are not normally connected to visual cortex pyramidal neurons {*2*}, indicating that the neuronal circuitry is better shaped to match endogenous patterns. Thus, also at this level integration improved in the *Trem2^-/-^* host environment. This is also in line with the molecular analysis using spatial transcriptomics. Importantly, tNs in *Trem2^-/-^* brains were transcriptionally highly correlated to endogenous layer II/III neurons, while tNs in the wt SWI environment transcriptionally clustered with astrocytes at 1mpt, suggesting the presence of a high degree of immature neuronal identity. All of these data thus demonstrate that the absence of TREM2 unlocks a checkpoint in tN functional maturation and integration.

Thus, TREM2-mediated signalling is key in restricting synapse and functional neuron maturation, and these results highlight the importance of regulating chronic microglia activation in neuronal transplantation. This is important to consider as clinical trials related to tNs focus so far largely on the identity of the neurons being transplanted, while not yet considering so much the adverse influence of the different pathological environments. Here we show that tN maturation and integration in acute brain injury would profit from reducing TREM2 activation, ideally dosing TREM2 signaling more precisely using recently established antibody tools {*61*}. This is important to consider, given that many attempts in neurodegenerative disease focus on activating TREM2 function, which should be avoided when neuronal replacement therapies are attempted. Our data suggest that creating a more permissive environment, specifically by attenuating TREM2-mediated chronic activation, may be as important as optimizing the neurons themselves

### Limitations of the study

Our findings provide novel insights into how different parameters of synaptic and functional integration may be misaligned for tNs in a pathological environment, highlighting the need to focus not only on a single read-out. Nevertheless, several limitations should be considered. We demonstrated that TREM2 plays a role in tNs integration and functional maturation, and it could now be interesting to proceed with cell-type-specific TREM2 deletion to pinpoint the exact cell type involved. Furthermore, it would be interesting to investigate whether, like other morphological, transcriptional and functional parameters, the ultrastructural synaptic features of tNs in *Trem2^-/-^*animals are also restored to endogenous levels. Finally, for therapeutic approaches using antibody tools that limit TREM2-mediated signaling may be preferable. This will bring us closer to the important aim of tN integration as close as possible to endogenous neurons thereby ensuring adequate functionality under all conditions.

## Materials & Methods

### Animals

Animal experiments were performed in compliance with German and European Union guidelines and approved by the Government of Upper Bavaria. All experiments were executed in the Core Facility Animal Models of the Biomedical Centre, LMU Munich. The animals were housed in specific pathogen–free conditions with a 12-hour light-dark cycle. Food and water were supplied *ad libitum*. Mice used for experiments were between 8 and 14 weeks of age at the time of the first surgery. The following mouse strains used: *Cx3cr1^GFP^* (B6.129P2(Cg)-Cx3cr1tm1Litt/J) [*62*), *Trem2^-/-^* (B6.129P2-TREM2tm1cln) {*63*}, β-*Actin-GFP* (C57BL/6-Tg(CAG-EGFP)1Osb/J) {*64*}, *Cux2-CreERT2* x *GFPrep* B6 ((Cg)-Cux2tm3.1(cre/ERT2)Mull/Mmmh x B6.Tg(CAG-eGFP)) MMRRC stock no.: 032779-MU and *C57bl6/J*.

### Surgeries

For all surgeries animals were anaesthetized using a mixture of fentanyl (0.05 mg/kg; Janssen), midazolam (5 mg/kg; Ratiopharm), and medetomidine (0.5 mg/kg; Orion Pharma). Once the animals lost their reflexes, the surgeries commenced. The head of the mouse was shaved, disinfected and the skin of the head was opened using a sterile scalpel. Topical 2% lidocaine was then applied to the skull prior to craniotomy. Once the surgery was completed, the skin was sutured (except for cranial window implantation, see below) and the anesthesia antagonized with atipamezol (2.5 mg/kg; Janssen), flumazenil (0.5 mg/kg; Hexal), and buprenorphine (0.1 mg/kg; Essex). Mice were placed on a heating pad until awake.

### Stab wound injury

Stab wound injury was performed as previously described (*48*) with minor modifications. Briefly, once anaesthetized, a craniotomy at the visual or somatosensory cortex was drilled. The skull piece was removed and stored in saline throughout the procedure. A 1mm-long, 0.5mm-deep stab wound injury was performed using an ophthalmological lancet with the following stereotactic coordinates from Lambda: 0.0 ± 0.3 antero-posterior and 2.0 ± 0.3 to 3.0 ± 0.3 medio-lateral for visual cortex and Bregma: −0.5 - -1.5 antero-posterior, −1.6 medio-lateral for somatosensory cortex. The skull piece was lightly dried from saline and placed to cover the cranial window. For the analysis of the stab wound only condition, the timepoint analysed was at 1-month post injury (mpi).

### Transplantation

Seven days after stab wound injury, donor cells were transplanted in the injured cortex as previously described {*7*}. Briefly, donor cells were either collected from embryonic day (e) 14.5-E15.5 (C57bl6/J or *Actin^GFP^*) cultures or acutely dissociated from e18 (*Actin^GFP^*) embryos. **e14.5 protocol**: Cultured cells were *in vitro* virally transduced using the viruses mentioned in the below section ‘Cortical neuron primary culture and viral labelling’. On the day of transplantation, cultured cells were washed three times using PBS, followed by trypsinization (Gibco, 25200-056), at a concentration of 0.025% for 10 minutes at 37°C. **e18 protocol**: On the day of surgery, cortical pieces from e18 embryos were dissected under a dissection microscope. They were then incubated with 0.025% Trypsin (Gibco, 25200-056) for 7 minutes at 37°C. **e14.5 & e18 protocol**: The trypsinized cells were then mixed with an FBS (PanBiotech, P30-3302) -containing medium in 1:1 ratio. After centrifugation, the cell pellet was resuspended in a medium containing high-glucose (4.5 g/l), GlutaMAX DMEM (Gibco, 61965-026), B27 (Gibco, 17504044) and penicillin-streptomycin (Gibco, 15140-122). A total volume of 1µl of the prepared cell suspension (50,000 – 100,000 cells/ µl) was transplanted using a 33-gauge Hamilton syringe at the following stereotactic coordinates from Lambda: 0.0 ± 0.3 antero-posterior, 2.5 ± 0.3 medio-lateral and 0.5-0.1mm dorso-ventral for visual cortex and Bregma: −1 antero-posterior, −1.6 medio-lateral and −0.6 – 0.1mm dorso-ventral for somatosensory cortex. After transplantation, the skull piece covering the cranial window was re-positioned. The timepoints analysed were at 1-month or 3-months post transplantation (mpt).

Adeno-associated virus injections

To analyze the spine density of endogenous neurons after stab wound injury in C57BL/6J mice, we performed adeno-associated virus (AAV) injections. To label endogenous neurons under stab wound injury conditions, we used a cocktail of PHP.B-capsid AAVs: pAAV-gfaABC1D-iCre (1×10⁹ GC), pAAV-CBh-FLEX-EGFP-W3SL (1×10¹⁰ GC), and pAAV-CBh-FLEX-DsRed2-W3SL (1×10¹⁰ GC) at a Cre virus : flip-excision switches virus ratio of 1:10. The AAVs were injected at 3-days post-injury (dpi), as shown in Fig. 1D. These constructs enabled mixed labeling of astrocytes and neurons. The AAV cocktail was injected into the stab wound site in the somatosensory cortex at a volume of 900 nL per location, delivered via an automated nanoinjector (Nanoliter, World Precision Instruments) at a rate of 40-60 nL/min. Mice were sacrificed at 1-month post-injection (mpi) for subsequent analysis.

### Rabies virus injections

To analyze whole brain connectivity of tNs in C57bl6J and *Trem2^-/-^* mice, retrograde monosynaptic RABV tracing was performed. RABV injection occurred at 4wpt (1mpt) or 12wpt (3mpt) as in Figure 5A. Depending on the construct expressed by tNs (see cell culture virus treatment section), the Rabies virus used was EnvA-pseudotyped SAD-ΔG-EGFP or SAD-ΔG-mCherry. To label endogenous neurons in the intact brain in 3-5-month old mice, we used a SAD-ΔG-EGFP, SAD-G pseudotyped RABV [*19*). This construct enables the RABV to infect preferentially neurons, due to the expression of the SAD glycoprotein in their envelopes. The RABV selected was injected in three locations surrounding the transplantation site in the visual cortex (for connectivity analysis) or in the intact visual cortex (endogenous neuron labelling), 200 nl per location using an automated nanoinjector (Nanoliter, World Precision Instruments) at 1nl/s. Mice were sacrificed 7 days after injection for subsequent analysis.

### Cranial window implantation surgery

For animals scheduled for live imaging experiments, a cranial window was implanted. After transplantation was performed, instead of returning the bone flap, a 4mm diameter cover glass (Warner instruments, 64-0724) was placed and sealed using histoacryl glue (B. Braun, 9381104) and dental cement (Pala Paladur, Kulzer; liquid, 64707937; powder, 64707945). This enabled the imaging of the transplanted cells in the injured visual cortex. Animals used for these experiments were either *Cx3cr1^GFP^* or *C57bl6/J*.

### Tamoxifen induction protocol

Adult Cux2-CreERT2 x GFPrep B6 were administered intraperitoneal Tamoxifen (Merck, 1.00983) at 40mg/kg twice daily for three days. Mice were sacrificed after five days for electron microscopy.

### Cortical neuron primary culture and viral labelling

The neocortex from e14.5 or e15.5 embryos was dissected and mechanically dissociated in HBSS (Gibco, 14025-050) with HEPES (Gibco, 14025-050). The cell suspension was then plated in a Poly-D-Lysine (PDL, Sigma, P0899) -coated, 24-well plate at a density of 200,000-300,000 cells per well and cultured in a medium containing high-glucose (4.5g/l), GlutaMAX DMEM, 10% FBS, 1% PenStrep. The cells were left to settle for 2-4 hours at 37°C and afterwards transduced with 1ul of Moloney murine leukemia virus-derived retroviral vectors. The following were the retroviral vectors used for the different experiments: connectivity analysis either CAG-DsRedExpress2-2A-Glyco-IRES2-TVA or CAG-TVA-2A-eGFP-2A-N2CG; electrophysiology - Syn-GFP; spine density, synapse, live imaging, Visium and electron microscopy (EM) analysis - CAG-GFP; live imaging - CAG-mScarlet. For the following experiments cortical cells from β-*Actin-GFP* were isolated and therefore no treatment with retroviruses was required: spine density, electrophysiology, EM, Visium HD. In the subsequent 2 days, the serum- containing medium was gradually replaced by a medium containing high-glucose, glutamax DMEM (4.5g/l), 1:50 B27, 1% PenStrep. Donor cells were left to culture for 3-5 days before transplantation.

### Tissue preparation

Mice were deeply anaesthetized and transcardially perfused using ice cold PBS and 1:200 Heparin (Braun, PZN:15782698), followed by 4% PFA (Carl Roth, 0335.4). The brains were then carefully dissected and post-fixed for 4-6 hours in 4% PFA. Afterwards, the brains were stored in PBS-Azide (G-Biosciences, 786-750). The brains were then either processed as free-floating sections or as cryosections. For free-floating section staining, the brains were sectioned sagittally 50-60µm thick using a vibratome (Leica VT1000S) and stored in PBS-azide. For cryosections, after post-fixation, the brains were incubated in 30% sucrose (Carl Roth, 4621.2) for 2 overnights at 4 °C. The brains were then embedded in NEG-50 (Epredia, 6502) and sectioned 20µm-thick sagittal sections using a cryostat (ThermoScientific CryoStar NX70). The sections were then stored at −70°C.

### Immunohistochemistry

The selected free-floating sections were first incubated with blocking solution (10% normal goat serum (Merck, S26), 300mM Glycine (Carl Roth, 3790.3), 0.2% Triton X100 (Carl Roth, 6683.1)) overnight at 4°C, followed by primary antibody incubation for 3 overnights at 4°C. Subsequently, sections were washed and incubated with secondary antibodies and Dapi (1:1000; Sigma, MBD0015) either overnight at 4°C or 2 hours at room temperature. In the case of synaptic staining, the sections were treated with citric acid for 30 minutes at 95°C prior to the blocking step.

The primary antibodies used are the following: anti-GFP (1:1000, Aves Lab, Cat# GFP-1020), anti-Iba1 (1:1000, Synaptic Systems, Cat# 234013), anti-P2RY12 (1:1000, AnaSpec Inc., Cat# AS-55043A). The secondary antibodies used were the following: Donkey anti-chicken 488 (1:500, Thermo Fisher Scientific, Cat#A78948), Donkey anti-rabbit 647 (1:500, Thermo Fisher Scientific, Cat#A31573), Goat anti-mouse IgG1 647 (1:500, Thermo Fisher Scientific, Cat#A21240), Goat anti-guinea pig 546 (1:500, Thermo Fisher Scientific, Cat#A11074), Donkey anti-rabbit Cy3 (1:500, Dianova, Cat#711-165-152).

#### *In situ* hybridization

*ln situ* hybridization of *Trem2* was performed using RNAscope® Multiplex Fluorescent Reagent Kit (ACD, 323110) according to manufacturer’s instructions. Briefly, 20 µm thick frozen brain sections were heated at 60°C for 30 minutes until completely dry. The sections were then fixed in 4% PFA and dehydrated in a graded ethanol series starting at 50% and ending with 100%. The sections were then treated with hydrogen peroxide at room temperature for 10 minutes and afterwards boiled in 1x target retrieval solution for 3 minutes, rinsed in Diethylpyrocarbonate (DEPC) water, followed by a short rinse in ethanol. After air dried, sections were treated with Protease III for 15 minutes at 40°C, rinsed in DEPC water, and incubated with the Trem2 probe (mm-Trem2-C3; 404111-C3) for 120 minutes at 40°C. Samples were rinsed in washing buffer and the signal amplified at 40°C using RNAScope Multiplex Fl v2 Amp1 for 30 minutes, RNAScope Multiplex Fl v2 Amp2 for 30 minutes, RNAScope Multiplex Fl v2 Amp3 for 15 minutes. Finally the signal was developed using RNAScope Multiplex Fl v2 HRP for 15 minutes at 40°C, followed by an incubation of the OPAL 570 (1:1000; Akoya Biosciences, FP1488001KT) for 30 minutes at 40°C, and ending this step with a 15 minute incubation with HRP-blocker at 40°C. Following this step, the immunohistochemistry protocol was followed.

### DNA-PAINT in brain tissue slices

#### Nanobody-DNA conjugation via single cysteine

Nanobodies against VGlut1 (1:150, Nanotag, Cat#N1605), PSD95 (1:150, Nanotag, Cat#N3705), Synaptotagmin1 (1:200, Nanotag, Cat#N2302), GFP (1:200, Nanotag, Cat#N0301), rabbit (1:200, Nanotag, Cat#N2405) and mouse kappa light chain (1:200, Nanotag, Cat#N1205) comprised of a single ectopic cysteine at the C-terminus for site-specific and quantitative conjugation. The conjugation to DNA-PAINT docking sites (see Table 3) was performed as described previously (*65*). First, the buffer was exchanged to 1x PBS and 5 mM EDTA, pH 7.0 using Amicon centrifugal filters (10k MWCO) and free cysteines were reacted with a 20-fold molar excess of bifunctional maleimide-DBCO linker (Sigma Aldrich, cat: 760668) for 2-3 hours on ice. The unreacted linker was removed by buffer exchange to PBS using Amicon centrifugal filters. Azide-functionalized DNA was added with 3-5 molar excess to the DBCO-nanobody and reacted overnight at 4°C. Unconjugated nanobody and free azide-DNA were removed by anion exchange using an AKTA Pure liquid chromatography system equipped with a Resource Q 1 ml column. Nanobody-DNA concentration was adjusted to 5 µM (in 1xPBS, 50% glycerol, 0.05% NaN3) and stored at −20°C.

**Table 3.**
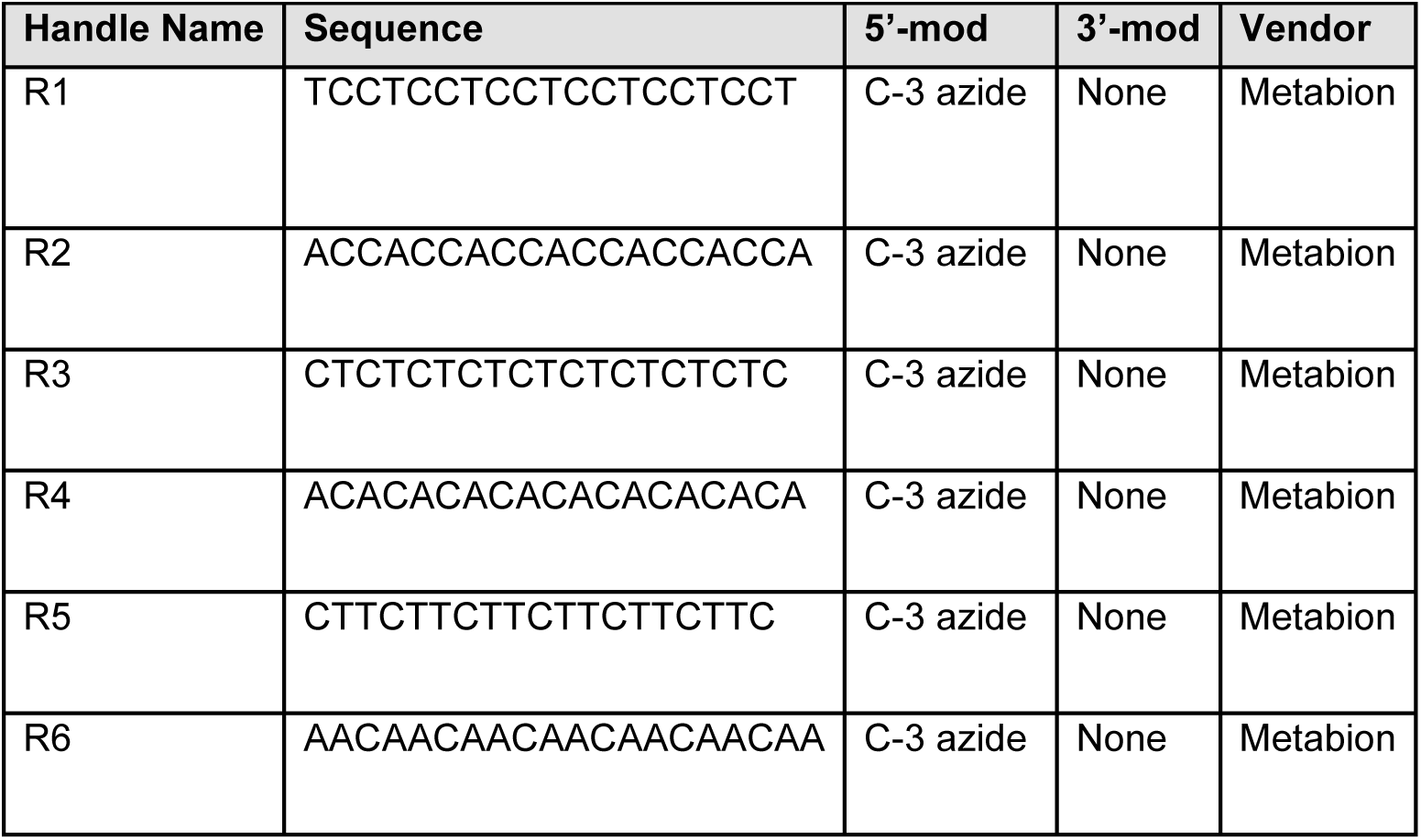
Docking Site Sequences used for the nanobody-DNA conjugation via a single cysteine.

| Handle Name | Sequence | 5'-mod | 3'-mod | Vendor |
| --- | --- | --- | --- | --- |
| R1 | TCCTCCTCCTCCTCCTCCT | C-3 azide | None | Metabion |
| R2 | ACCACCACCACCACCACCA | C-3 azide | None | Metabion |
| R3 | CTCTCTCTCTCTCTCTCTC | C-3 azide | None | Metabion |
| R4 | ACACACACACACACACACA | C-3 azide | None | Metabion |
| R5 | CTTCTTCTTCTTCTTCTTC | C-3 azide | None | Metabion |
| R6 | AACAACAACAACAACAACAA | C-3 azide | None | Metabion |

#### DNA-PAINT microscopy setup

Fluorescence imaging was carried out on a custom-built setup using an inverted microscope (Nikon Instruments, Eclipse Ti2) equipped with the Perfect Focus System, using an objective-type TIRF configuration equipped with an oil-immersion objective (Nikon Instruments, Apo SRHP TIRFx100, NA 1.49, Oil). A 561 nm laser (MPB Communications, 1 W) is used for excitation and coupled into the microscope via a Nikon manual TIRF module. The laser beam is passed through a cleanup filter (Chroma Technology, ZET488/10x for 488 nm excitation and ZET561/10× for 560 nm excitation) and coupled into the microscope objective using a beam splitter (Chroma Technology, ZT561rdc). Fluorescence light is spectrally filtered with an emission filter (Chroma Technology, ET600/50m, and ET575lp) and imaged on an sCMOS camera (Hamamatsu Fusion BT) without further magnification, resulting in an effective pixel size of 130 nm after 2x2 binning, leading to an imaging field of view of approximately 67x67 µm. Highly inclined and laminated optical sheet (HiLO) (*66*) illumination was used for all measurements. For imaging a central region of interest of 1,152 × 1,152 pixels (576 × 576 after binning) of the camera were chosen. 3D imaging was performed using a cylindrical lens (Nikon Instruments, N-STORM) in the detection path. The camera readout sensitivity is set to 16-bit and the readout bandwidth to 200 MHz. The scan mode of the camera was set to “ultra quiet scan” (readout noise = 0.7 e-r.m.s., 80 μs readout time per line). Image acquisition and microscope control is performed using µManager (Version 2.0.1) {*67*}.

#### 6-plex DNA-PAINT imaging

Round glass coverslips (18 mm, Th.Geyer Labsolute, cat: 9161062) were PDL-coated (Sigma, P0899) and stored at 4 °C until further use. For DNA-PAINT imaging, a coated coverslip was incubated with 90 nm gold nanoparticles for 10 min to serve as fiducials, followed by four vigorous washes with 1xPBS. A single tissue slice was then immobilized by drying on the coated coverslip. Permeabilization was performed for 24 h at 4 °C using 1% Triton in 1xPBS. The tissue was subsequently washed three times with PBS, and blocking buffer (1x PBS, 3% BSA (Sigma Aldrich, A4503-10G), 1% Triton X-100 and 0.05 mg/ml sheared salmon sperm DNA (Thermo Fisher Scientific, 15632011)) was applied for at least 2 h at 4 °C. After blocking, the sample was again washed three times with 1xPBS before immunolabeling reagents were applied in antibody incubation buffer (1xPBS, 1 mM EDTA, 0.02% Tween-20, 0.05% NaN_3_, 2% BSA, and 0.05 mg/ml sheared salmon sperm DNA, 1% Triton) and incubated overnight at 4 °C. The nanobodies conjugated above and the following antibodies against Gephyrin (1:200, Synaptic Systems, Cat#147 011) and vGAT (1:200, Invitrogen, Cat#PA5-27569) were used for imaging.

The following morning, the sample was washed four times with 1xPBS and once with buffer C wash (1x PBS, 0.1 mM EDTA, 500 mM NaCl; pH 7.4). The coverslip was then mounted onto a custom-made sample chamber, fixed in place using two-component silicone glue, to enable buffer exchange between imaging rounds. The first imaging solution in buffer C+ (1x PBS, 0.1 mM EDTA, 500 mM NaCl and 0.05% Tween-20; pH 7.4) together with 1x PCD, 1x PCA, and 1x Trolox, was applied to the sample according to Table 4. 100x Trolox were prepared by: 100 mg Trolox was added to 430 μl 100 % Methanol, 345 μl 1M NaOH, and 3.2 ml H_2_O. 40x PCA were prepared by: 154 mg PCA, 10 ml water and NaOH were mixed and pH was adjusted 9.0. 100x PCD were prepared by: 9.3 mg PCD, 13.3 ml of buffer (100 mM Tris-HCl pH 8, 50 mM KCl, 1 mM EDTA, 50 % Glycerol). Suitable transplanted regions for imaging were identified using the 488 nm laser line. All targets were imaged using the 560 nm laser line at 85 W/cm^2^ laser power at the objective in HILO illumination mode, acquiring 20.000 frames at 100 ms exposure time (except for GFP, where 30.000 frames were acquired). Between targets the sample was washed with at least 2 ml buffer C wash until no residual signal from the previous imager solution was detectable. The next imaging solution was then introduced and this process was repeated until all six targets were imaged.

**Table 4.**
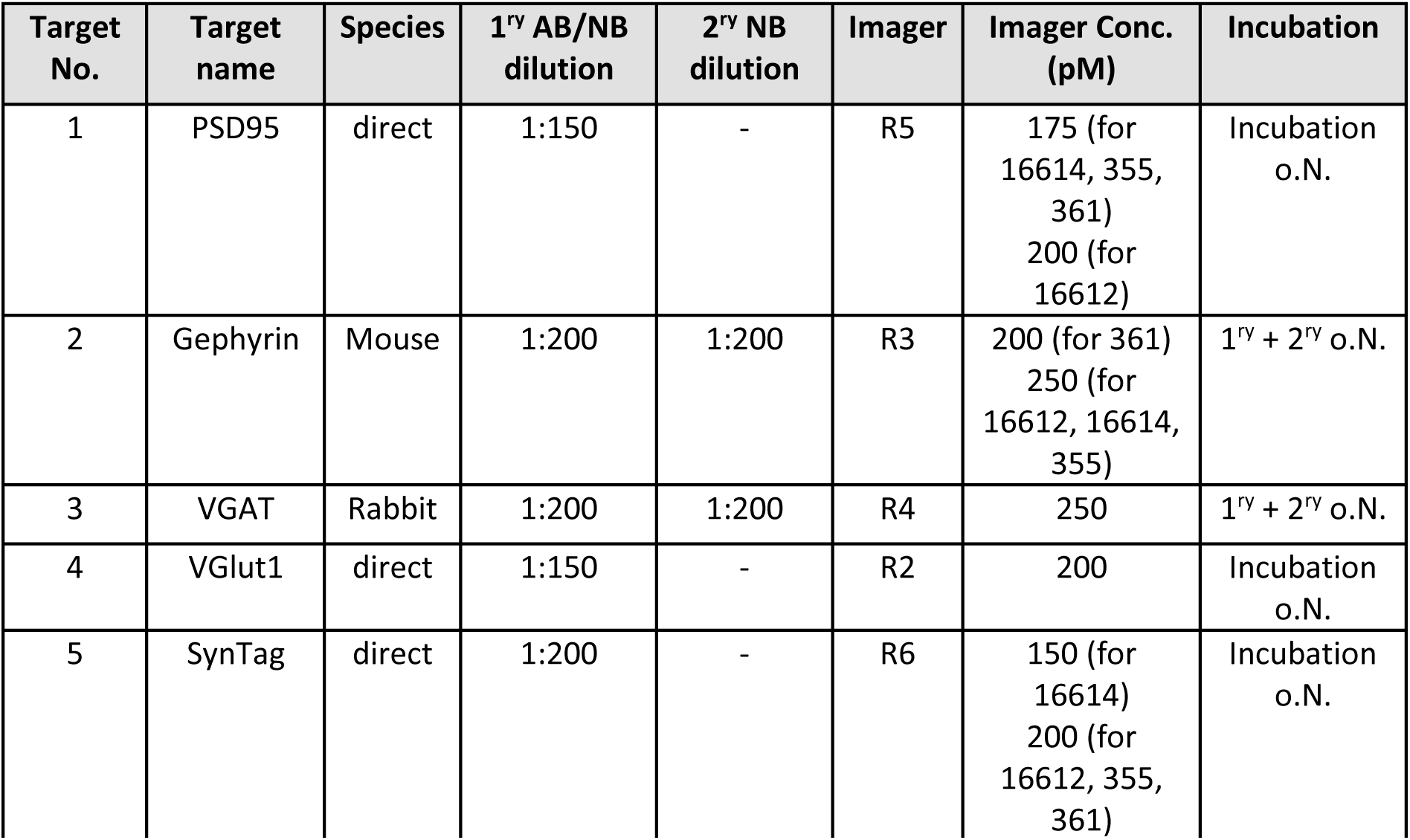
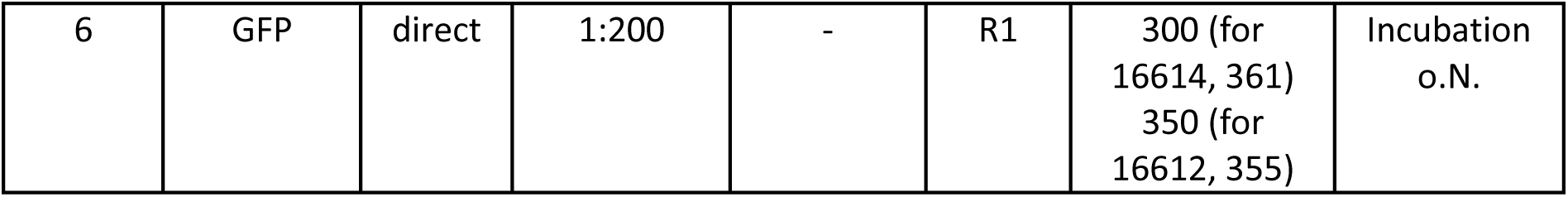
Experimental parameters for DNA-PAINT imaging.

### 2-photon live imaging

For the chronic longitudinal *in vivo* imaging of transplanted cells in the injured brain, cranial windows were implanted on the day of transplantation as described above. For the initial microglia morphology, the mice were imaged at 1, 2, 3 and 4wpt, whereas for the longitudinal dendritic length quantification, the mice were imaged at 4, 5, 6 wpt for 1mpt and 12, 13wpt for the 3mpt timepoints. For each imaging session, mice were anaesthetized using a mixture of fentanyl (0.05 mg/kg; Janssen), midazolam (5 mg/kg; Ratiopharm), and medetomidine (0.5 mg/kg; Orion Pharma) and subsequently placed on a custom-designed, head-fixing device, which fitted onto the microscope imaging stage. tNs were imaged at 15Hz with an Olympus MPE-RS resonant scanner multi-photon system using a femto-second pulsed Ti:Sapphire laser. The laser was tuned at 840nm for eGFP and 1040nm for mScarlett detection. A 25x water-immersion Olympus objective with a 1.25 numerical aperture was used, with the following imaging settings: z-spacing −0.50mm, pixel resolution – 1024, dwell time - 2-6ms. After an imaging session, and the anesthesia was antagonized with atipamezol (2.5 mg/kg; Janssen), flumazenil (0.5 mg/kg; Hexal), and buprenorphine (0.1 mg/kg; Essex). After the final imaging session, the mice were sacrificed. Mice which developed cloudy cranial windows were excluded from the study.

### Electron microscopy (EM)

Animals were deeply anaesthetized and transcardially perfused with HBSS followed by 2% paraformaldehyde (Electron Microscopy Sciences (EMS), 15710), 2% glutaraldehyde (EMS, 16220) in PBS. The fixative was prepared freshly and used within 2 hours. All solutions were warmed to 37°C immediately prior to use. The brains were carefully dissected and post-fixed overnight at 4°C. The following day, the brains were sectioned at 50µm using a vibratome (Leica VT1000S). Sections were subsequently fixed on ice for an additional 1 hour and afterwards washed with PBS. The region of interest was identified using an epifluorescent microscope (Zeiss Imager M2) and dissected under a stereomicroscope (Olympus S2X10). The sections were processed for EM max. 24h later for optimal tissue preservation and restoration of antigenicity. Slices were transferred to fixative solution and incubated for 1h on ice.

The slices were washed in 0.1M PB (Soerensen buffer, EMS, 11601-10) on ice and transferred into 50 mM glycine (Sigma) for 30 min, followed by another washing step in 0.1M PB. Sucrose infiltration was started with 15% (1-2 h) and 30% (overnight) sucrose each while gentle agitation at 4°C. The tissue slices were placed onto a folded piece of aluminum foil with the slice side facing down and excessive sucrose solution wiped using a clean disposable wipe. The slice is hovered over liquid nitrogen using forceps until slices and residual sucrose turn rigid and white, then the foil is dipped into liquid nitrogen for 1 min. The sample is taken out and warmed up to room temperature until slices become transparent again. After a washing step in 0.1M PB the slices detach from the aluminium foil be gentle pipetting solution onto them. The slices are transferred into TBS (Tris buffered saline, Sigma-Aldrich, T5912) for washing and blocked in 10% normal goat serum (Aurion, 900.077) and 1% fish skin gelatin (Aurion, 900.033) in TBS for 1h at 4°C with gentle agitation. The primary antibody in diluted (1:10) blocking solution (anti-GFP (abcam ab6556), 1:5000) was incubated for 2d at 4°C with gentle agitation. After extensive washing in TBS the nanogold conjugated secondary antibody (nanogold IgG-GAR nanoprobes, 1:100, in diluted (1:10) blocking solution) was incubated for 1d at 4° C with gentle agitation. The slices were washed in PBS, post-fixed in 1% glutaraldehyde in PBS for 10 min and thoroughly washed with water. For silver enhancement we used the kit (HQ Silver, nanoprobes, #2012-45ml) according to the manufacturer’s protocol. Briefly, two drops of solution A and two drops of solution B were mixed in a round bottom 2 mL microcentrifuge tube, vortexed and two drops of solution C added, followed by vortexing. One slice was transferred immediately after mixing using a brush, submerged and incubated for 5 min in the dark. The reaction was stopped by addition of water and the coloring of the GFP-positive region evaluated under a dissecting microscope. The reaction was fully stopped in 0.1M PB and slices stored over night at 4° C in 0.1 M PB.

The slices were washed in water and then dried out shortly before addition of a few drops followed by 1.5 mL 1% osmium tetroxide (EMS) and 1.5% potassium ferrocyanide (Sigma) solution in 0.1 M sodium cacodylate (Science Services) buffer (pH 7.4) for 1h at 4° C. The staining was enhanced by reaction with 1% thiocarbohydrazide (Sigma) for 20 min at 40 °C, followed by washing and another incubation in 1% osmium tetroxide for 30 min. The slices were immersed into 1% aqueous uranyl acetate (Science Services) at for 1h at 4°C in the dark, washed and dehydrated in in an ascending ethanol series. The resin LX112 (LADD) was infiltrated in acetone and blocks were cured for 2d a 60°C and trimmed (TRIM2, Leica). Serial sections, with a nominal thickness of 50 nm, were generated using a 35° ultra-diamond knife (Diatome) on an ATUMtome (Powertome, RMC). These sections were collected onto carbon-coated Kapton tape (kindly provided by Jeff Lichtman and Richard Schalek), freshly treated with plasma (custom-built, based on Pelco PELCO easiGlow, adopted from M. Terasaki, U. Connecticut, CT). The tape stripes were affixed onto adhesive carbon tape (Science Services), and this assembly was then secured onto 4-inch silicon wafers (Siegert Wafer), and grounded through adhesive carbon tape stripes (Science Services). EM micrographs were acquired using an Apreo S2 SEM (Thermo Fisher Scientific) as previously described (*68*). Hierarchical imaging of serial sections was achieved through initial mapping of the plastic tape stripes at low lateral resolution and capture of the entire tissue sections at a medium resolution (ranging from 100 nm to 200 nm). Correlation of the region of interest was established through the integration of vasculature and slice morphology. Subsequently, micrographs of serial sections were acquired at a lateral resolution of 3 nm. To align the serial section imaging data, a hybrid approach combining both automated and manual processing procedures was employed within Fiji TrakEM2 [*69*). Image segmentation was performed in VAST {*70*}.

### Electrophysiology

Mice were briefly anaesthetized with isoflurane and rapidly killed by decapitation. Coronal brain sections (150-200μm thick) containing the transplanted cortical area were cut in an ice-cold high-sucrose, low-sodium artificial cerebral spinal fluid (ACSF) and then maintained in normal ACSF at 37°C for 30–45min. Afterwards sections were stored in a slice-maintenance chamber at room temperature (∼22°C) until transfer into the recording chamber. Composition of the normal ACSF in mM: NaCl 125, KCl 2.5, NaHCO3 26, glucose 10, NaH2PO4 1.25, sodium pyruvate 2, myo-inositol 3, CaCl2 2, MgCl2 1, and ascorbic acid 0.5 pH was 7.4, bubbled with 95% O2, 5% CO2. For the low-sodium ACSF CaCl2 and MgCl2 concentrations were 0.1 and 4mM, respectively, and NaCl was replaced by 200mM sucrose. Experiments were conducted at 36 ± 1°C, maintained by an inline feedback temperature controller and heated stage (Warner Instruments) with the recording chamber being continuously perfused with ACSF at a rate of 1–2 ml min−1. Whole-cell patch-clamp recordings were made from visually identified V1 neurons using an EPC10/2 HEKA amplifier (HEKA Elektronik), sampled at 50 kHz and filtered at 10 kHz. Patch pipettes were pulled from borosilicate glass capillaries (Warner Instruments) using using a 2-stage vertical puller (PC-10 Narishige, Tokyo, Japan), filled with a patch solution containing (in mM): K-gluconate 126, KCl 4, HEPES 40, EGTA 5 MgCl2 1, Na2phosphocreatine 5, 0.2% biocytin, 292mOsm. pH was adjusted to 7.2 with KOH. Data were corrected for liquid junction potentials of –13.8mV. Electrode resistance was between 2.4 and 6 MΩ. Spontaneous action potential firing was recorded at the neurons’ resting membrane potential. Spontaneous excitatory synaptic inputs were recorded at a holding potential of −70mV. Based on our internal pipette solution, the calculated chloride reversal potential was very close to the resting membrane potential. To increase the IPSC driving force and enable better assessment of IPSCs, they were recorded at membrane potentials of −50mV, slightly more depolarized, but still somewhat physiological. Synaptic responses were evoked by afferent fiber stimulation with concentric bipolar electrodes (FHC inc., #CBARC75). Voltage pulses were generated by the HEKA amplifier and post-amplified by an isolated pulse stimulator (AM Systems). Neuronal properties were exported from Patchmaster (HEKA Elektronik) and analyzed using Clampfit software (Molecular Devices). n is the number of neurons, with 2-3 brain slices per animal and at least 3 animals per group.

### Viral vector cloning and production

The AAV plasmids, pAAV-CBh-FLEX-EGFP-W3SL and pAAV-CBh-FLEX-DsRed2-W3SL were cloned based on the in-house pAAV-CBh-FLEX-MCS-W3SL vector, inserting GFP via fragment ligation with NheI/EcoRI and DsRed2 with NheI/BamHI-Blunt. pAAV-gfaABC1D-iCre was cloned on vector pAAV-gfaABC1D-tdTomato (was a gift from Baljit Khakh, addgene 44332) {*71*} replacing tdTomato with codon improved Cre (was a gift from Jinhyun Kim, addgene 51904) {*72*} via PCR with BamHI-Blunt/MfeI. pRV-Syn-GFP was cloned based on pRV-CAG-GFP, replacing CAG with the Synapsin 1 promoter via PCR with Blunt-XhoI. pRV-CAG-GFP, pRV-CAG-mScarlet, pRV-CAG-TVA-2A-eGFP-2A-N2cG were cloned via Gateway cloning. In brief, the destination plasmids pRV-CAG-DEST were originally from pLenti7.3/V5-DEST™ Gateway™ Vector (Life Technologies) and exchanged to the MML-RV backbone, and pENTR1A-TVA-2A-eGFP-2A-N2cG (was a gift from Ian Wickersham, addgene 100811) was cloned via Gibson assembly. All viral plasmids were produced as endotoxin-free plasmids using either a CsCl density gradient or an endotoxin-free kit (Qiagen, 12362). RV and rAAV vectors were produced and titered as described in {*73*}.

### Circuit mapping processing

#### Whole brain staining and clearing

After perfusion, brains were post-fixed for 24 hours at 4°C and finally 1 hour at room temperature. Afterwards stored in PBS-Azide. For brain staining and clearing, the nanobody(V_H_H)-boosted 3D imaging of solvent-cleared organs (vDISCO) method was used {*74*}^69^. Briefly, whole brains were incubated for 2 days at 37°C in pre-treatment solution (1.5% goat serum, 0.5% Triton X100, 0.5mM Methyl-beta-cyclodextrin (Sigma-Aldrich, 332615), 0.2% trans-1-acetyl-4-hydroxy-L-proline (Sigma-Aldrich, 441562), 0.05% PBS-sodium azide). Followed by 10-14days at 37°C in fresh pre-treatment solution, 1:400 FluoTag X4 anti-GFP Alexa 647 (NanoTag; N0304-AF647), and FluoTag X2 anti-mScarlet-I AZDye568 (NanoTag; N1302-AF568) or FluoTag-X2 anti-mScarlet Atto 565 (cat no. N1302-At565) nanobodies and 1:1000 Dapi. After staining, the brains were washed 4 times using a wash solution (1.5% goat serum, 0.5% Triton X100, 0.05% PBS-sodium azide), followed 4 PBS washes at room temperature. Dehydration of samples was performed using different steps of tetrahydrofluran (THF, Sigma-Aldrich, 186562) of 50%, 70%, 80% and 100% of 2 hours each at room temperature, with the final step repeated with an overnight incubation at room temperature. For clearing, the brains were incubated for 1 hour in 100% THF, followed by 1 hours in Dichloromethane (DCM; Sigma-Aldrich, 270997) and finally stored in 1 part benzyl alcohol (BA; Sigma-Aldrich, 402834) and 2 parts benzyl benzoate (BB; Sigma-Aldrich, B6630). The brains were left at least 1 overnight in BABB solution, for equilibrium of refractive index to be reached.

#### Imaging of the cleared samples with light-sheet microscopy

We used a 4x objective lens (Olympus XLFLUOR 340) equipped with an immersion corrected dipping cap mounted on a LaVision UltraII microscope coupled to a white light laser module (NKT SuperK Extreme EXW-12) set at a constant power level at 95% for imaging. The images were taken in 16-bit depth at a nominal resolution of 1.625 μm/voxel on the XY axes. Gross anatomy was scanned via tissue autofluorescence using a 470/30 excitation filter and a 535/50 emission filter. The visualization of the ATTO555/AZDye568 used a 580/25 excitation and a 625/30 emission filter and Alexa 647 was acquired using a 640/40 excitation and a 690/50 emission filter. In z-dimension we took the sectional images in 4 μm steps using left and right sided illumination. The nominal thinnest point of the light-sheet was 4.06 μm using a sheet NA of 0.090. All samples were scanned with a rectangular 2x3 tiling. The reconstruction of the datasets from the tiling volumes was performed by stitching the acquired multichannel tiled volumes using TeraStitcher’s automatic global optimization function (v1.11.1).

### Spatial transcriptomics

#### Tissue processing &sequencing

Adult C57Bl6/J mouse brains were used for spatial transcriptomics using Visium protocol as recommended by 10x Genomics and as previously described (*75*, *76*). **For Visium experiment:** For the control brain sections only a stab wound injury was performed and sacrificed at 1-month post injury (mpi; *n* = 1; Fig. 4A). For the transplantation brain tissue tN were injected 1 week after stab wound injury and sacrificed at 1mpt (*n* = 2; Fig. 4A) as previously published in (*30*]. **For the Visium HD experiment:** one mouse per condition was used to analyse wt versus *Trem2^-/-^* at 1mpt and 3mpt (4 mice; n=1).*^-^*The brains were snap frozen in an isopentane (Carl Roth #6752.5) and liquid nitrogen bath and stored at −80°C. The frozen brains were sectioned at 10 µm thickness using a cryostat (Thermo Scientific, Cryostar NX70] and resected to comprise of only the cortical layers, corpus callosum and hippocampal brain regions. This allowed the fitting of 2 brain sections per capture area. **For Visium experiment:** The tissue was stained using H&E staining and imaged with the Carl Zeiss Axio Imager.M2m Microscope using a 10x objective. The libraries were prepared using the Visium Spatial Gene Expression Reagent Kits (10x Genomics PN-1000187, protocol: <u>CG000239</u>) with 18 min permeabilization time. Sequencing was performed on an Illumina HiSeq1500 or NovaSeq 6000 System, and a paired-end flowcell (SP lane Dual Indexed Sequencing Run). Sequencing was performed at the Helmholtz MUnchen Genomics Core Facility. Data were mapped against the mouse reference genome mm10 (GENCODE vM23/Ensembl 98; builds versions 1.2.0 and 2020A from 10x Genomics) with Space Ranger 1.2.2.). **For the Visium HD experiment:** The sections were first stained for H&E according to the 10x Genomics guidelines and imaged with an AxioScan 7 digital slide scanner (Zeiss, Oberkochen,Germany) equipped with a 20x magnification objective. A CytAssist machine was used in the Visium HD experiment to transfer the HE stained section on the donor slide to the Visium HD slide. The staining and the downstream processing of the Visium HD samples were processed according to manufacturer’s protocol (10X Genomics CG000684 | Rev A). The generated libraries were sequenced using Nextseq P2 flowcell.

### Image Analysis

#### Dendrite length quantification

To monitor the dendrite lengths of transplanted cells from the longitudinal live imaging sessions, images from 4, 5 or 6 wpt were quantified for 1mpt and for the 3mpt imaging timepoints 12 or 13 wpt were analyzed. Image quality determined the selection for analysis. Two-photon images were processed using Fiji (ImageJ){*77*}. Raw images were converted from .oir files to .tif files using the plugin OlympusViewer. For neurons processes spanning more than one field of view, adjacent images were taken and subsequently stitched. Stitching was done using the program ImarisStitcher 9.6.0 (Bitplane). Dendrite lengths were quantified using the Fiji plugin ‘Simple Neurite Tracer’.

#### Spine density & type quantification

Confocal images of dendrites from transplanted neurons were obtained with an Olympus FV1000 using a 60x UPlanSApo oil objective (NA 1.35) or a Leica SP8 Falcon with a 63x HC PL APO CS2 objective (NA = 1.4). Images were acquired using the following settings: pixel resolution 800 x 800, z-step of 0.2µm,

6x zoom and a scan speed of 8us/pixel (Olympus) or pixel size of 0.037µm, z-step of 0.2µm, 6x zoom and a scan speed of 200Hz (Leica). Transplants from e14.5 cultured cells were transduced with a CAG-GFP virus, whereas e18.5 transplanted cells were acutely dissociated from *Actin^GFP^*-positive embryos. Raw images were deconvoluted with Huygens Essential v.16.05 (Scientific Volume Imaging). Spine density and type were quantified using the ImageJ (Fiji) {*77*} plugin ‘Cell counter’. Spines were defined similarly as before {*78*}, which were any protrusions emanating from the dendritic shaft, and preferentially lateral. Spine types were defined based on morphology. Mushroom spines had a large head with respect to their thin neck (Fig. 1E), filopodia were any spines longer than 1µm (Fig. 1F), thin spines were spines that their head was as thin as the neck (Fig. S1F), stubby spines were protrusions which did not possess a neck (Fig. S1I), branched spines had visibly two heads but joined into one stalk before reaching the shaft (Fig. S1F), and spine head filopodia were like mushroom spines with an additional small protrusion emanating from the head (Fig. S1). Dendrite length was measured using the plugin ‘Simple Neurite Tracer’ {*79*}.

#### DNA-PAINT Single-molecule localization analysis

Raw fluorescence data were reconstructed using the Picasso software package {*80*} (the latest version is available at https://github.com/jungmannlab/picasso). Drift correction was performed using the AIM algorithm {*81*}^77^ with gold nanoparticles as fiducials for all experiments. Alignment between different channels was done through cross-correlation of the fiducial gold nanoparticles using Picasso.

Clusters of synaptic proteins were obtained using the DBSCAN algorithm {*82*} and minimum localizations and clustering radius were selected based on the imaging parameters of the individual super resolution channel of the protein. To define a minimum localization threshold, background regions were analyzed and used as a baseline. Both DBSCAN parameters were further adjusted based on visual inspection to determine a cutoff value that distinguishes background noise from specific protein clusters.

Synapse were manually segmented using the Picasso {*80*} pick tool with a circular selection region of 7 pixels (910 nm). A clearly identifiable signal of the postsynaptic scaffolding proteins PSD95 and gephyrin were used as selection criteria. For synapses located at the transplant site, regions were selected based on the presence of postsynaptic scaffolding proteins within a maximum distance of 500 nm from the GFP signal marking the transplant. Cluster volumes for the postsynaptic scaffold proteins are calculated by finding a 3D convex hull and extracting its volume. For presynaptic proteins VGAT, VGlut1, and Synaptotagmin1, only clusters with more than 200 localizations were considered. Localization counts above this threshold were extracted from all selected synaptic regions. Synapses were subsequently classified into 8 different groups according to their pre- and postsynaptic protein composition. Classification criteria used were as follows: Excitatory chemical synapses (PSD95, vGLUT1 and/or SynTag), PSD95 empty (PSD95 only), inhibitory chemical synapse (gephyrin, vGAT and/or SynTag), gephyrin empty (gephyrin only), and mixed synapse (gephyrin and vGlut1). The identified synapse classes were used for plotting synapse ratios on GFP-positive dendrites. The excitatory synapse population consisted of excitatory chemical synapses and PSD95 emtpy, whereas the inhibitory synapse population consisted of inhibitory chemical synapses, gephyrin empty, and mixed synapses. Synapses with a mixture of synaptic proteins, not previously described, were omitted from the analysis.

#### Trem2 in situ quantification

Cryosections stained with *in situ* hybridization for *Trem2* were used to validate the expression of *Trem2* at the transplantation site. Confocal images of the transplantation/injury site (intratransplant/intralesion) and a nearby field of view without the presence of tNs/injury (peritransplant/perilesion) were obtained using a Leica SP8 Falcon with a 20x HC PL APO IMM CORR CS2 objective (NA = 0.75). The images were acquired using the following settings: pixel size of 0.541µm, z-step of 1.043µm, and a scan speed of 200Hz. Using ImageJ (Fiji), the images were collapsed to their maximum intensity projection and a region of interest (ROI) based on the tN fluorescence was traced. The mean gray intensity of *Trem2* was measured. The same ROI was measured for the corresponding peritransplant image. Normalized values represent the mean gray intensities values obtained from intratransplant image divided by those from the corresponding peritransplant image.

#### Microglia morphology quantification

Cryosections stained for Iba1 labelling microglia were used for this analysis. Confocal images of the transplantation site (intratransplant) and a nearby field of view without the presence of tNs (peritransplant) were obtained using a Leica SP8 Falcon with a HC PL APO 20x IMM CORR CS2 (NA = 0.75). The images were acquired using the following settings: zoom 2x, pixel size of 0.271µm, z-step of 0.514µm, and a scan speed of 200Hz. The microglia morphology was analysed using Imaris 9.6.0 (Bitplane) using the ‘filament’ function. Normalized values represent the microglial filament measurements obtained from intratransplant images divided by those from the corresponding peritransplant image.

#### Microglia sphericity quantification

Images from the 2-photon live imaging (above) from 1-4wpt were used. Microglia were assessed by the CX3CR1 fluorescence. These images were analysed using Imaris 9.6.0 (Bitplane) using the ‘surfaces’ function

#### Starter cell quantification

After whole-brain imaging, cleared brains were glued to a glass dish and immersed in BABB. Two-photon images of the transplanted area were acquired using a Leica SP8 Falcon DIVE with an upright BABB-resistant 16x HC FLUORTAR L IMM CORR VISIR dip-in objective (NA = 0.6). The images were acquired with 1024x1024 pixels, pixel size of 0.339µm, z-step of 1.625µm and a scan speed of 200Hz (lines/second). Image stacks of different colors were recorded stack sequential to minimize lengthy laser wavelength adjustments (∼20 s). This led to slight drift in xy between the colors which was corrected in Imaris 9.6.0. Afterwards the images were manually analysed using the cell counter function in ImageJ (Fiji). The number of starter cells was quantified based on a neuronal appearance and the co-localization of the two fluorophores stemming from the transduction of tNs and the infection of RABV.

#### Presynaptic cell quantification

Labeled neurons were detected using the cellfinder software as previously described [*83*). Briefly, the detection process consisted of three main steps. First, potential cell candidates were identified using three-dimensional filtering and thresholding techniques. Instead of the default global thresholding method provided by cellfinder, we employed a local thresholding approach to account for the decreased brightness of labeled cells deeper within the brain compared to those near the surface. Second, the identified cell candidates were automatically classified as either “cells” or “non-cells” using a 3D adaptation of the ResNet convolutional neural network {*84*}, which had been pre-trained on a subset of the data. Finally, the classification results were manually reviewed and corrected, if necessary, using a napari-based graphical user interface {*85*} integrated within cellfinder to ensure accurate detection. After manual inspection, the data were registered to the Allen Mouse Brain Atlas {*86*} using the registration pipeline implemented in cellfinder {*87*}.

#### EM annotation & synapse quantification

EM images were manually segmented and annotated using VAST 1.4.0. The 1mpt and 3mpt datasets were processed as above, whereas the used wt dataset is a published dataset from (*25*]. For each condition, at least 5 dendrites were analysed. tN were identified based on their morphology and the GFP-immuno gold dark spots present at their soma. Synapses were identified based on the presence of pre- and postsynapse structures and the presence of vesicles at the presynapse. The identification of symmetrical synapses and asymmetrical synapses was performed as previously described (*26*]. Briefly, in addition to the synapse criteria, the presence (asymmetrical synapses) or absence (symmetrical synapses) of a postsynaptic density was assessed across the z-slices. Skeleton annotations were extracted using excel. Dendrites shorter than 15µm were omitted from quantification.

### Bioinformatic analysis

For Visium: Analysis of the spatial datasets was followed according to the Seurat protocol {*88*}. Briefly, the spatial transcriptome datasets were converted to Seurat objects, merged and integrated using the Harmony batch effect correction algorithm {*31*}, recommended by 10X Genomics. Followed by a dimensionality reduction and unsupervised clustering, visualized in UMAP space. To assess the cell composition and upregulated GOterms, the significant (adjusted p value <0.05, >1.5 log2FC) transcripts in cluster 16 were assessed using CellMarker 2024 {*32*} and GOrilla analysis [*89*] respectively. To assess the presence of microglial-specific sensome markers, the top 25 genes from [NO_PRINTED_FORM] {*33*}, Supplementary table 2 were downloaded and this list was compared to the significant transcripts in cluster 16. To assess the presence of TREM2-associated microglia signature, the top 10 genes mostly upregulated in DAM stage 2 vs. Stage 1 signature from {*34*}, Supplementary table 7 were downloaded and a module score created.

For Visium HD: All libraries were aligned to a custom reference genome containing GRCm39 and GFP sequence and probe reference using spaceranger. For the *Trem2^-/-^* 3mpt sample, manual image alignment was done in the 10X Genomics Loupe Browser v9.0.0.. The datasets were then loaded and processed in python using squidpy and spatialdata and afterwards brough to single cell resolution using bin2cell. Cells were kept if they had between 200 and 8000 counts, a minimum of 100 genes, and less than 25% mitochondrial reads. Genes were kept if they were present in at least 5 cells. For visualization, the scanpy spatial functions were used. Clustering was done using the leiden algorithm. Clusters were then annotated based on their spatial location and marker gene expression from the Allen Brain Atlas. GFP-positive cells were identified based on whether they had at least 1 GFP read. Differential expression analysis between the final cell types was done using sc.tl.rank_genes_groups after removing cell types that were present in only one of the datasets.

### Statistical analysis

Statistical details of experiment including statistical test used, level of statistical significance and number of animals/cells/dendrites can be found in the figure legends. Microsoft excel was used to compile the data and the statistical significance was analysed using GraphPad Prism 7. For data acquired with patch-clamp recording; Statistical analyses of the data were performed with SigmaStat/SigmaPlot. Prior to every statistical test a Shapiro-Wilk normality test was run to determine if a parametric or non-parametric test was to be used. For samples where normal distribution could be confirmed the following tests were used: for two samples comparison, unpaired two-tailed t test; for more than 2 samples, one-way ANOVA followed by Tukey’s multiple comparison test. In non-normally distributed samples we used: for two samples comparison nonparametric, Mann-Whitney test and Wilcoxon test; for more than 2 samples Kruskal-Wallis followed by Dunn’s multiple comparison test. To correlate 2 parameters a linear regression analysis was performed and when more than one variable need to be examined a 2-way ANOVA followed by Tuckey’s multiple comparison test was performed. Obtained p values were stated as significance levels in the figure legends and marked in the graphs by asterisks (***p < 0.0005; **p < 0.005; *p < 0.05, ns > 0.05) and all data are presented as means ± SEM.

## Resource Availability

Requests for further information, resources, and reagents are available upon further request

## Acknowledgements

We are grateful for the excellent technical assistance of T. Simon-Ebert and D. Franzen and the great discussions about synaptic integration of neurons in adult neurogenesis with Armen Saghatelyan. We thank the Mesoscale Hub of SyNergy including the expertise of Ali ErtUrk. We also thank the BMC Bioimaging facility and the expertise of Steffen Dietzel. Multi-photon microscopy on fixed tissue was performed at the Core Facility Bioimaging of the Biomedical Center, funding for the Leica SP8 MP WLL Falcon microscope was provided by the Deutsche Forschungsgemeinschaft, grant number INST 86/1909-1. For the EM experiments, we would like to acknowledge the contribution of Cornelia Niemann, SyNergy and TRR274 funding for the Nanoscale Hub. We acknowledge the technical support of Genomics Core Facility and Pathology & Tissue Analytics Core Facility and at Helmholtz Munich. We thank Inti Alberto De La Rosa Velazquez, Annette Feuchtinger, Richard Lindner, Melisa Gomez, Ulrike Buchholz, Theresa Isermenger for support with the Visium HD experiment. This study was supported by the German Research Foundation TRR274 (Nr. 408885537, M.G.), FOR2879/2 (Nr. 405358801 to M.G.), the Munich Cluster for Systems Neurology (EXC 2145 SyNergy-ID 390857198 to M.G., J.N., C.H. and R.P.) as well as by the EU in the consortium NSC Reconstruct (H2020, Nr. 874758) and the advanced ERC grant (Nr. 885382) to M.G. Parts of this work were supported by a New Frontiers in Research Fund Transformation grant to M.G., funded through three Canadian federal funding agencies (CIHR, NSERC, and SSHRC). Y.Z. has received funding from the European Union’s Horizon 2020 research and innovation programme under the Marie Skłodowska-Curie (grant agreement No 101024862). M.F.M-R. was supported by a Boehringer Ingelheim Fonds (BIF) PhD fellowship.

## Author Contribution

M.G. and Y.Z. conceived, designed, and co-ordinated the project. Y.Z. performed all surgeries except for electrophysiology (M.T.), spatial RNA-seq (Visium HD – M.T., and 1mpsw - C.K., V.S.), 1mpi with AAVs (C.L.), and 1mpi (M.T.). Y.Z. conducted all experiments except for spatial RNA-seq: Visium (M.F.M-R (1mpt), C.K. and V.S. (1mpi)) and Visium HD (Helmholtz Facility), electrophysiology (O.M.), whole brain imaging (Mih.T.), RNAscope (M.T.), DNA-PAINT (E.M.S.) and electron microscopy (G.K. and M.S.). Y.Z. analysed all data except for presynaptic partner analysis (E.P), DNA-paint (E.M.S.), Visium HD (M.R.) and electrophysiology (C.K-S.). E.P. and R.P. devised the semi-automated quantification of presynaptic partners. A.K. and M.K. assisted, provided expertise and instruments for the execution of 2-photon live imaging experiments. M.W. and C.H. provided mice and expertise on TREM2 and microglia. M.F.M-R helped with setting up the protocol for immuno-EM. M.P. helped with setting up the protocol for cleared brain staining. K.K.C. provided expertise and viral vectors for monosynaptic tracing. M.G. and Y.Z. wrote the manuscript with input from all co-authors.

