## Supplemental Fig. 1-6 for "Multimodal synapse analysis reveals limitations in transplanted neuron integration mediated by TREM2"

### Supplementary figure legends

**Supplementary figure 1. Spine density and type quantification of tNs after SWI. (A)** Spine density quantification of tNs at 1mpt and 3mpt. The transplanted cells were acutely dissociated from e18.5 embryos and directly transplanted (mice (n) = 4-5, dendrites = 42 - 80). **(B)** Spine density quantification of tNs of e14.5 cultured cortical cells and e18.5 acutely dissociated cortical cells. Data for e14.5 tNs were taken from Figure 1F (mice (n) = 4-5, dendrites = 42 - 80)**. (C-F)** Quantifications and their respective representative confocal images of 1mpt, 3mpt and endogenous neurons of **(C)** branched, **(D)** thin, **(F)** spine head filopodia and **(F)** stubby spine types. (mice (n) = 5, dendrites = 53 - 75)**.** In graphs, solid colors represent per-animal means; lighter shades represent individual measurements, with distinct shades denoting each mouse. All data are presented as means ± SEM. All statistics were performed on the means of individual mice - ns p >0.05, * p <0.05, ** p <0.005, *** p <0.0005 by unpaired two-tailed t-test (A), one-way ANOVA with Tukey's multiple comparison test (B-F).

**Supplementary figure 2. Ultrastructural analysis of tNs using ATUM-SEM. (A)** Scheme of brain tissue processing for electron microscopy analysis. Brain sections containing transplantation site are selected (left), transplantation site resected for immune-EM processing (blue box) and 3D EM dataset acquired (red box). 3D reconstructed tN neuron from EM image stack (green box) and a zoom-in of segment depicting tN (green) and synapses (orange). Shaft synapses are marked with blue arrows and spine synapses are labelled with magenta arrows. **(B-D)** Representative image of immunogold-labelled **(B)** tNs, **(C)** neurons in *Cux2-Cre x GFP* brain sections and **(D)** microglia in *CX3CR1*$^{GFP}$. GFP immunogold-labelled cells are indicated using green arrowheads and dendrite in (C) is indicated with orange arrowheads. **(E)** Representative sequential images from EM immunogold-labelling. tN dendrite is pseudo-colored in cyan. Empty spines (green), together with spine (magenta), and shaft (blue) synapses are indicated in arrows. **(F-G)** Graphical representations illustrating the distribution of the different synaptic parameter in control, 1mpt, and 3mpt. Graphs depict the percentage synapse composition at the **(F)** synapse, and **(G)** dendrite level. **(H-I)** Graphical representations illustrating the distribution of the different synaptic parameter in 1mpt according to different neurons analyzed. Graphs depict the percentage synapse composition at the **(H)** synapse, and **(I)** dendrite level. **(J)** Representative sequential images from EM immunogold-labelling. Asymmetrical synapses (cyan), and symmetrical synapses (orange) are indicated in arrows. All data are presented as means ± SEM. ns p >0.05, * p <0.05, ** p <0.005, *** p <0.0005 by two-way ANOVA (H, I).

**Supplementary figure 3. Electrophysiological properties of tNs (A)** Patch-clamp recording of spontaneous activity of tNs and control neurons in acute brain slices. Pie charts demonstrating the percentage of silent neurons. **(B-D)** Quantifications of **(B)** input resistance, **(C)** membrane constant, and **(D)** resting membrane potential in tNs and control neurons. Input resistance (Ri) was quantified by applying 50 repetitions of small-amplitude (10 pA) current steps, and calculating Ri from the average steady-state voltage deflection using Ohm's law. Decay tau was assessed from fitting an exponential decay function to the voltage defection. Resting membrane potential was recorded in the absence of current injection. **(E-G)** Afferent synaptic stimulation of eEPSCs in tNs in acute brain slices. **(E)** Typical examples of eEPSCs in control (black), 1mpt (orange) and 3mpt (blue) neurons. **(F)** Stimulation of disynaptic excitatory inputs were found in each experimental group. Stimulus artifacts have been omitted from the traces. **(G)** Pie charts of percentage of neurons that exhibited eEPSCs in response to afferent stimulation. All data are presented as means ± SEM. Data obtained from different mice is denoted in different shades. ns p >0.05, * p <0.05, ** p <0.005, *** p <0.0005 by One-way ANOVA test (C, D), Kruskal-Wallis test (B).

**Supplementary figure 4. Presence of reactive microglia around tNs expressing elevated levels of *Trem2*. (A)** The spatial information of the 18 clusters observed. **(B)** The spatial information of cluster 16, indicating its enrichment around the transplantation site (Red). SWI (left) and transplantation (right) site are highlighted with a yellow dotted line. **(C)** Scheme of the experimental setup used for live imaging between 1wpt and 4wpt. **(D)** Representative 2-photon images of 1, 2, 3 and 4 wpt of tNs (magenta) and the surrounding microglia (cyan). **(E)** Graphical representation of microglia sphericity quantification between 1-4wpt. Solid colors represent per-animal means; lighter shades represent individual measurements (mice (n) = 4, microglia = 193-684). **(F)** Scheme of the experimental setup used for live imaging between 1mpt and 3mpt. **(G)** Representative 2-photon images of peritransplant (top) and intratransplant (bottom) regions. tNs (magenta) and the surrounding microglia (cyan). Insets contain representative microglial cells in the region. **(H)** Representative confocal images of 1mpt (top) and 3mpt (bottom) of wt (left) and *Trem2*$^{-/-}$ (right) animals depicting microglia (Iba1; magenta) and tNs (GFP; green). **(I-L)** Graphical representation of quantified microglial features from immunohistochemical images in **H** at the transplantation site normalized to the values obtained from peritransplant regions. The graphs illustrate average microglia **(I)** filament length, **(J)** segments, **(K)** branch points, and **(L)** terminal points. **(M)** Images showing the expression of *Trem2* (magenta) around the transplantation site at 3mpt. Transplanted cells express GFP (green). **(N)** Graph comparing the normalized mean gray values of *Trem2* expression at 1mpt, and 3mpt. Data for 1mpt were taken from Figure 4L (mice = 4). All data are presented as means ± SEM. All statistics were performed on the means of individual mice - ns p >0.05, * p <0.05, ** p <0.005, *** p <0.0005 by repeated measure one-way ANOVA with Tuckey's multiple comparison test (E), One-way ANOVA test with Tuckey's multiple comparison test (I-L), and Mann-Whitney test (N).

**Supplementary figure 5. TREM2 is a tN circuit remodeler. (A)** Dot plot of the average expression (dot color) and percentage expressed (dot size) of the TREM2-associated microglial signature in 1mpsw and 1mpt spatial transcriptomics. **(B)** UMAP plot showing that the TREM2-associated microglial signature is concentrated in cluster 16. **(C-E)** Graphical representation of the connectivity ratios in wt and *Trem2*$^{-/-}$ at 1mpt (orange) and 3mpt (blue) in **(C)** ipsilateral sub-cortical areas, **(D)** contralateral cortical areas, **(E)** ipsilateral cortical areas. **(F)** Graphical representation of the correlation of number of presynaptic partners in the visual cortex and number of starter cells at 1mpt (orange) and 3mpt (blue) in *Trem2*$^{-/-}$ (linear regression, $R^2$ = 0.2632, p>0.05). **(G)** Color-coded graphical representation of the connectivity ratios of tNs in wt and *Trem2*$^{-/-}$ in the different brain regions at 1 and 3mpt (mice (n) = 3-4). **(H-M)** Graphical representations of tNs' spine type at 1mpt and 3mpt in *Trem2*$^{-/-}$ animals. The spine types analysed were **(H)** mushroom, **(I)** filopodia, **(J)** thin, **(K)** stubby, **(L)** branched, and **(M)** spine head filopodia. (mice (n) = 4, dendrites = 31 - 37). In graphs, solid colors represent per-animal means; lighter shades represent individual measurements, with distinct shades denoting each mouse. All data are presented as means ± SEM. All statistics were performed on the means of individual mice - ns p >0.05, * p <0.05, ** p <0.005, *** p <0.0005 by unpaired two-tailed t-test (H-M).

**Supplementary figure 6. Spatial transcriptome analysis reveals reduced inflammatory signature and increased tN synaptic maturation in a *Trem2*$^{-/-}$ environment. (A)** UMAP analysis of unsupervised clustering showing the four datasets overlapped without requiring batch-effect correction. **(B)** Spatial plots overlaying the cluster identities associated with GFP signal, for the different conditions. The presence of identified GFP spots is delineated with a dotted blue line, marking the transplantation site.

**(C)** Significantly top10 upregulated biological processes based on the DEGs in the transplant cluster in wt and *Trem2*$^{-/-}$ at 1mpt. **(D)** The top 10 significantly upregulated WikiPathways upregulated in the transplant cluster in wt brains. **(E)** Highlighted transcripts (green – both 1 and 3mpt; blue – 1mpt, orange – 3mpt) detected in our spatial dataset in the wt involved in the DAP12 microglial causal network. Network from WikiPathways. **(F)** Combined correlation matrix of the annotated cell types in wt and *Trem2*$^{-/-}$ at 1mpt and 3mpt. Individual matrices separated by timepoints are depicted in Fig. 7F, G.

### Supplementary Figure 1.

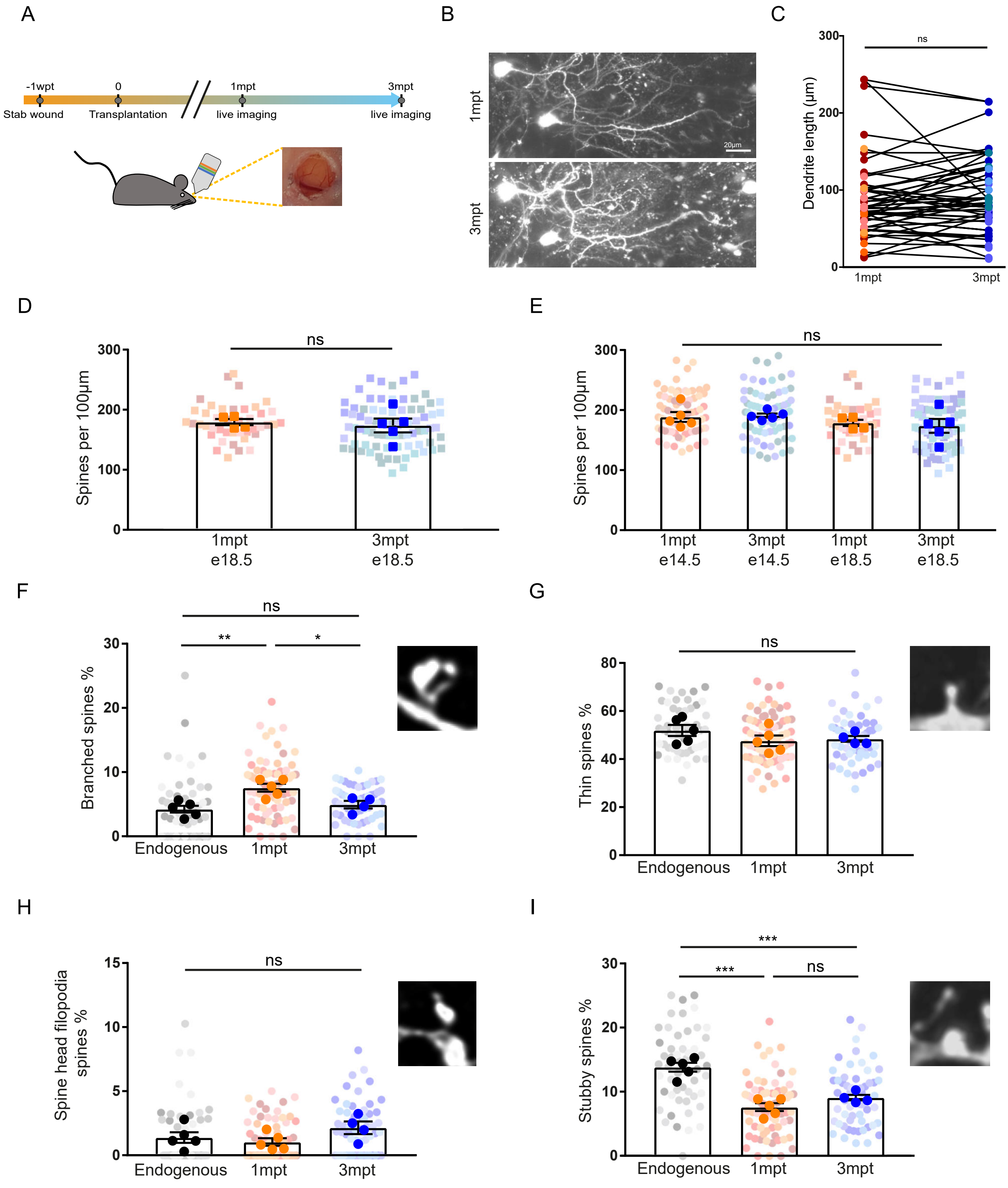

### Supplementary Figure 2.

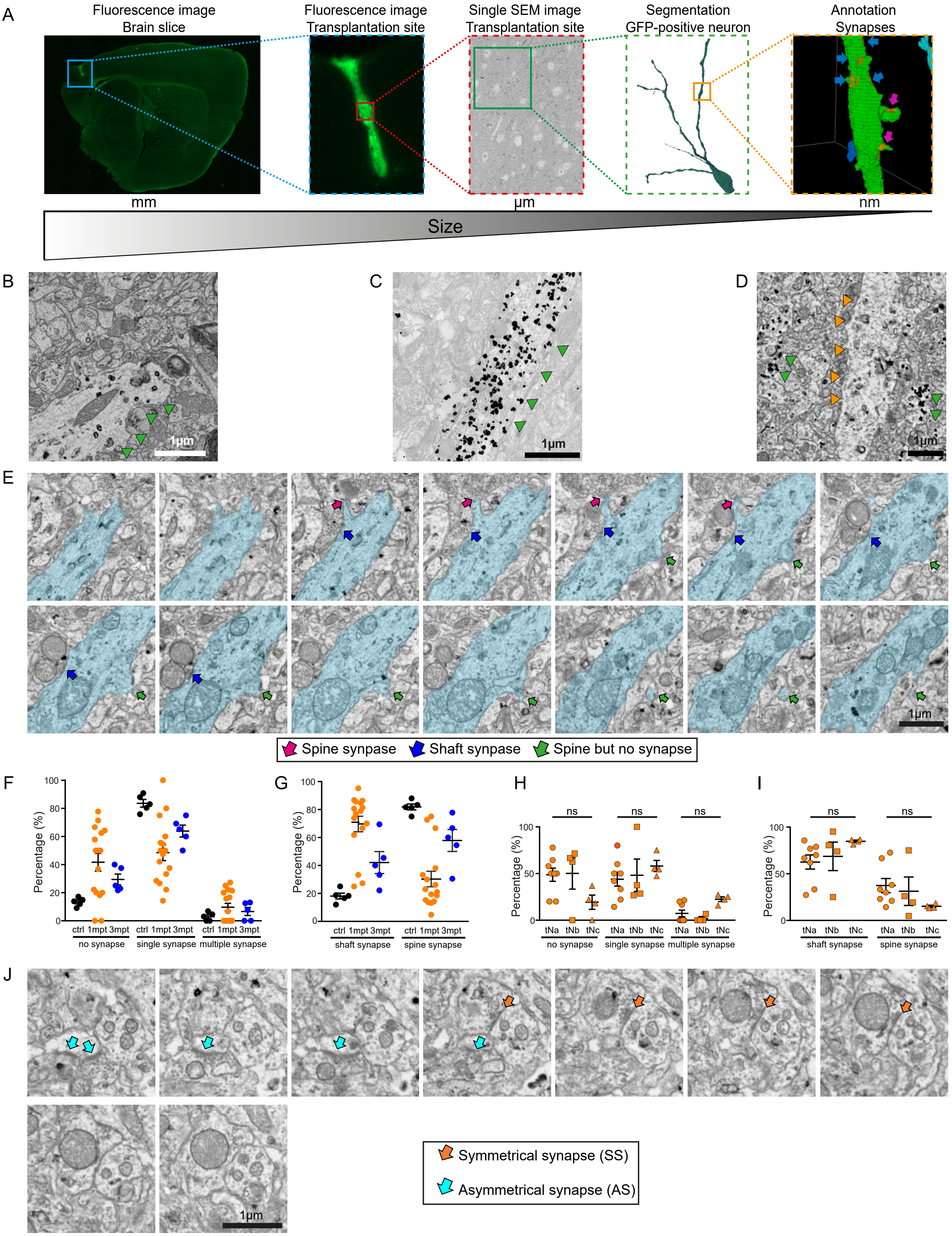

### Supplementary Figure 3.

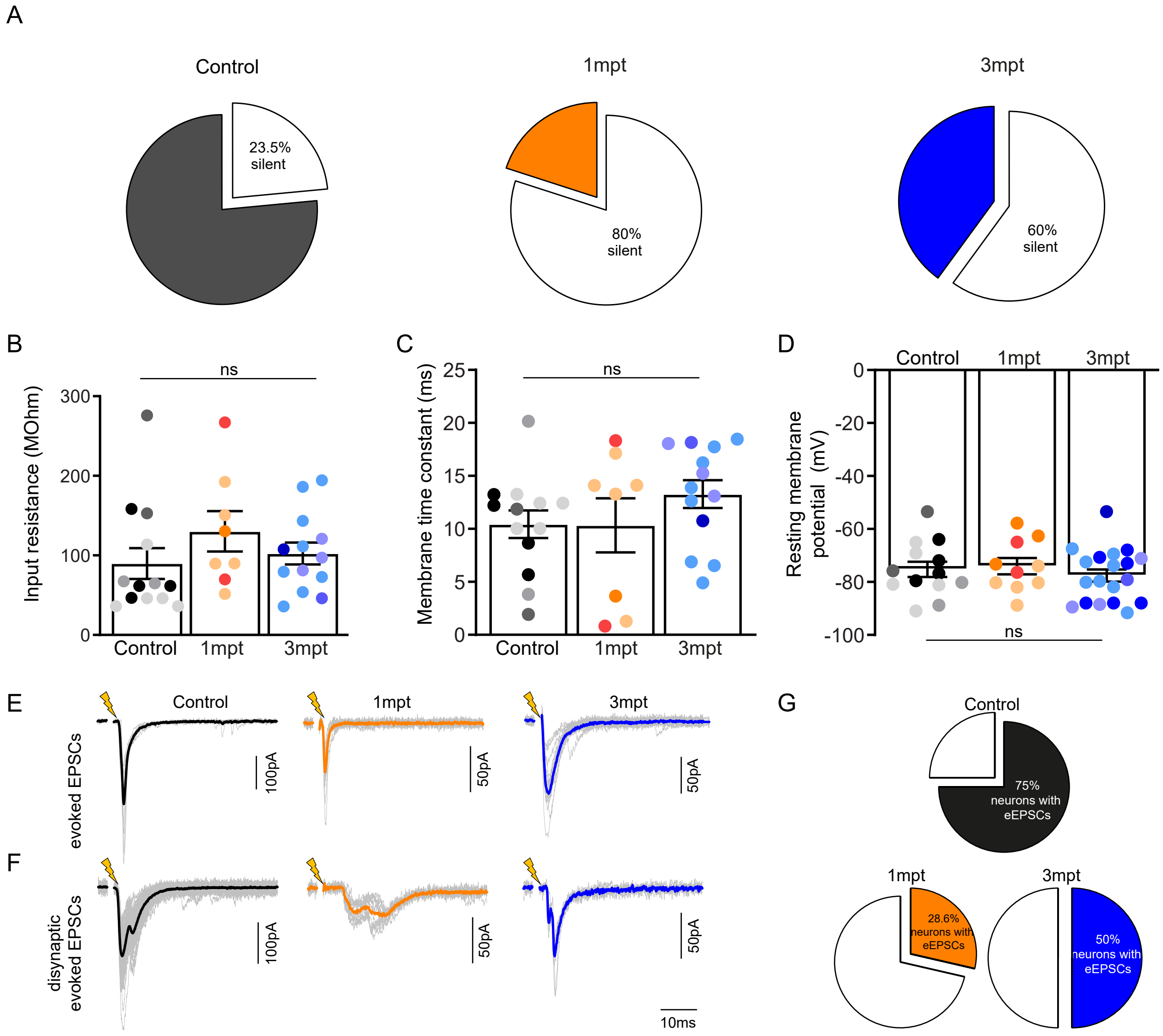

### Supplementary Figure 4.

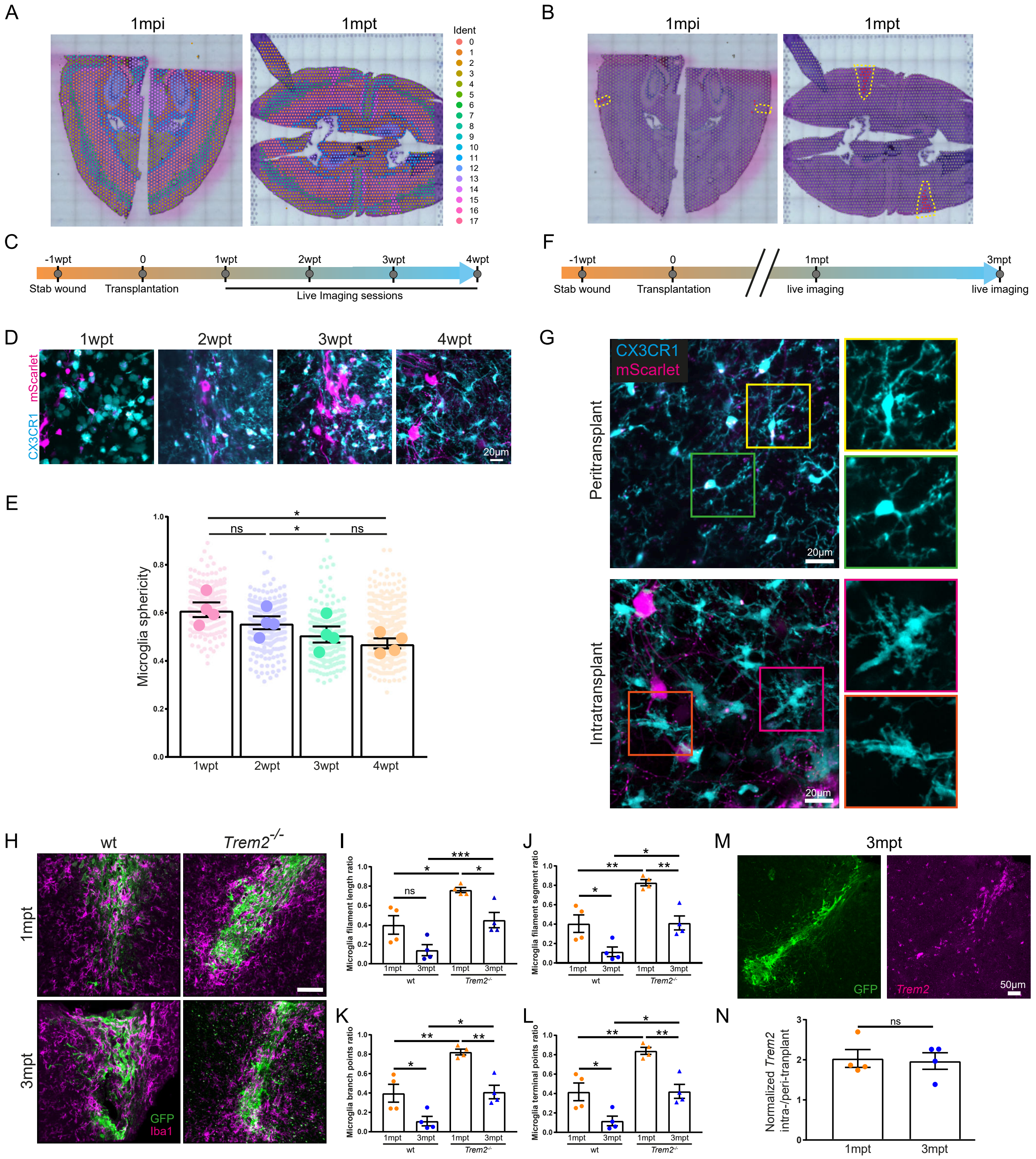

### Supplementary Figure 5.

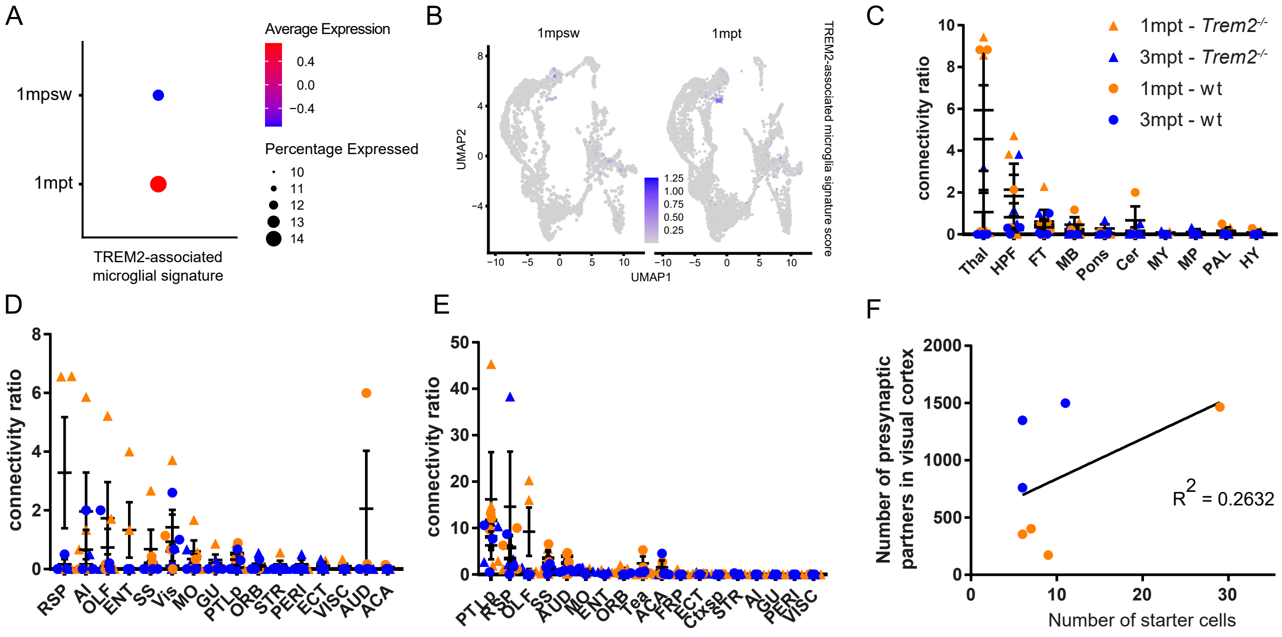


G

>100.00 | 50.00 - 99.99 | 5.00 - 49.99 | 1.00 - 14.99 | 0-01 - 0.99 | 0

| wt | | 1mpt | 3mpt |
|---|---|---|---|
| IPSI | Vis | 63.86 | 37.30 |
| | PTLp | 11.64 | 6.26 |
| | RSP | 5.82 | 3.46 |
| | AUD | 3.92 | 0.77 |
| | SS | 3.52 | 1.28 |
| | Tea | 2.39 | 0.52 |
| | ACA | 0.94 | 1.57 |
| Cortical | MO | 0.51 | 0.54 |
| | OLF | 0.06 | 0.00 |
| | AI | 0.00 | 0.00 |
| | ORB | 0.11 | 0.11 |
| | ECT | 0.12 | 0.00 |
| | Ctxsp | 0.05 | 0.00 |
| | STR | 0.00 | 0.11 |
| | VISC | 0.05 | 0.00 |
| IPSI | Thal | 4.77 | 0.00 |
| | | 0.43 | 0.33 |
| | HPF | 0.83 | 0.21 |
| Sub-cortical | MB | 0.46 | 0.00 |
| | Cer | 0.67 | 0.00 |
| | PAL | 0.17 | 0.00 |
| | HY | 0.10 | 0.00 |
| | Pons | 0.04 | 0.00 |
| Contra | Vis | 0.60 | 1.42 |
| | PTLp | 0.55 | 0.32 |
| | RSP | 0.00 | 0.17 |
| | AUD | 2.06 | 0.00 |
| | SS | 0.22 | 0.00 |
| Cortical | Tea | 0.00 | 0.00 |
| | ACA | 0.05 | 0.00 |
| | MO | 0.11 | 0.00 |
| | OLF | 0.06 | 0.73 |
| | AI | 0.00 | 0.50 |
| | ORB | 0.04 | 0.11 |

| *Trem2*-/- | | 1mpt | 3mpt |
|---|---|---|---|
| IPSI | Vis | 44.03 | 162.59 |
| | PTLp | 16.20 | 8.12 |
| | RSP | 1.48 | 14.62 |
| | OLF | 9.24 | 0.59 |
| | SS | 2.45 | 1.57 |
| | AUD | 1.19 | 2.49 |
| | MO | 0.64 | 0.90 |
| | ENT | 0.41 | 0.49 |
| Cortical | ORB | 0.82 | 0.00 |
| | Tea | 0.19 | 0.81 |
| | ACA | 0.82 | 0.03 |
| | FRP | 0.31 | 0.00 |
| | ECT | 0.09 | 0.25 |
| | Ctxsp | 0.06 | 0.06 |
| | STR | 0.06 | 0.06 |
| | AI | 0.04 | 0.00 |
| | GU | 0.00 | 0.06 |
| | PERI | 0.03 | 0.00 |
| IPSI | Thal | 4.55 | 1.06 |
| | HPF | 2.14 | 1.83 |
| | FT | 0.61 | 0.44 |
| | MB | 0.25 | 0.15 |
| Sub-cortical | Pons | 0.00 | 0.28 |
| | Cer | 0.03 | 0.17 |
| | MY | 0.06 | 0.06 |
| | MP | 0.04 | 0.11 |
| | PAL | 0.08 | 0.00 |
| | HY | 0.04 | 0.00 |
| Contra | RSP | 3.28 | 0.00 |
| | AI | 1.96 | 0.17 |
| | OLF | 1.73 | 0.00 |
| | ENT | 1.33 | 0.00 |
| | SS | 0.68 | 0.14 |
| | Vis | 0.93 | 0.11 |
| | MO | 0.60 | 0.34 |
| | GU | 0.31 | 0.20 |
| Cortical | PTLp | 0.49 | 0.06 |
| | ORB | 0.04 | 0.14 |
| | STR | 0.14 | 0.00 |
| | PERI | 0.06 | 0.17 |
| | ECT | 0.07 | 0.11 |
| | VISC | 0.08 | 0.00 |
| | AUD | 0.08 | 0.00 |

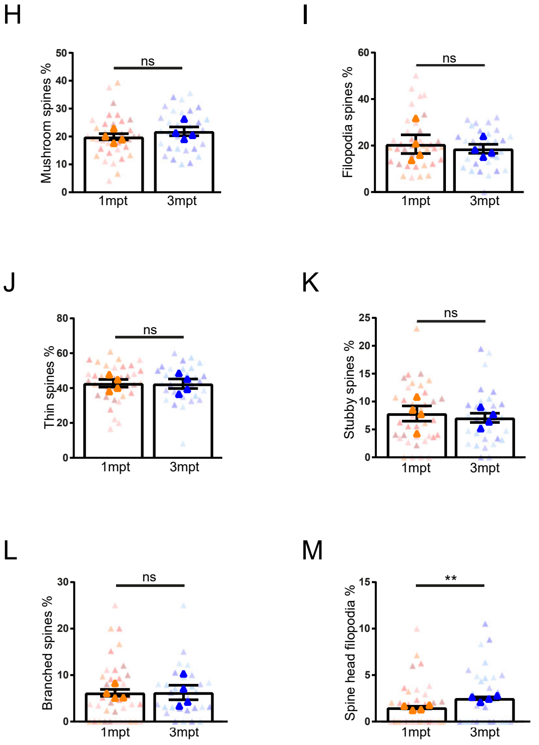

### Supplementary Figure 6.

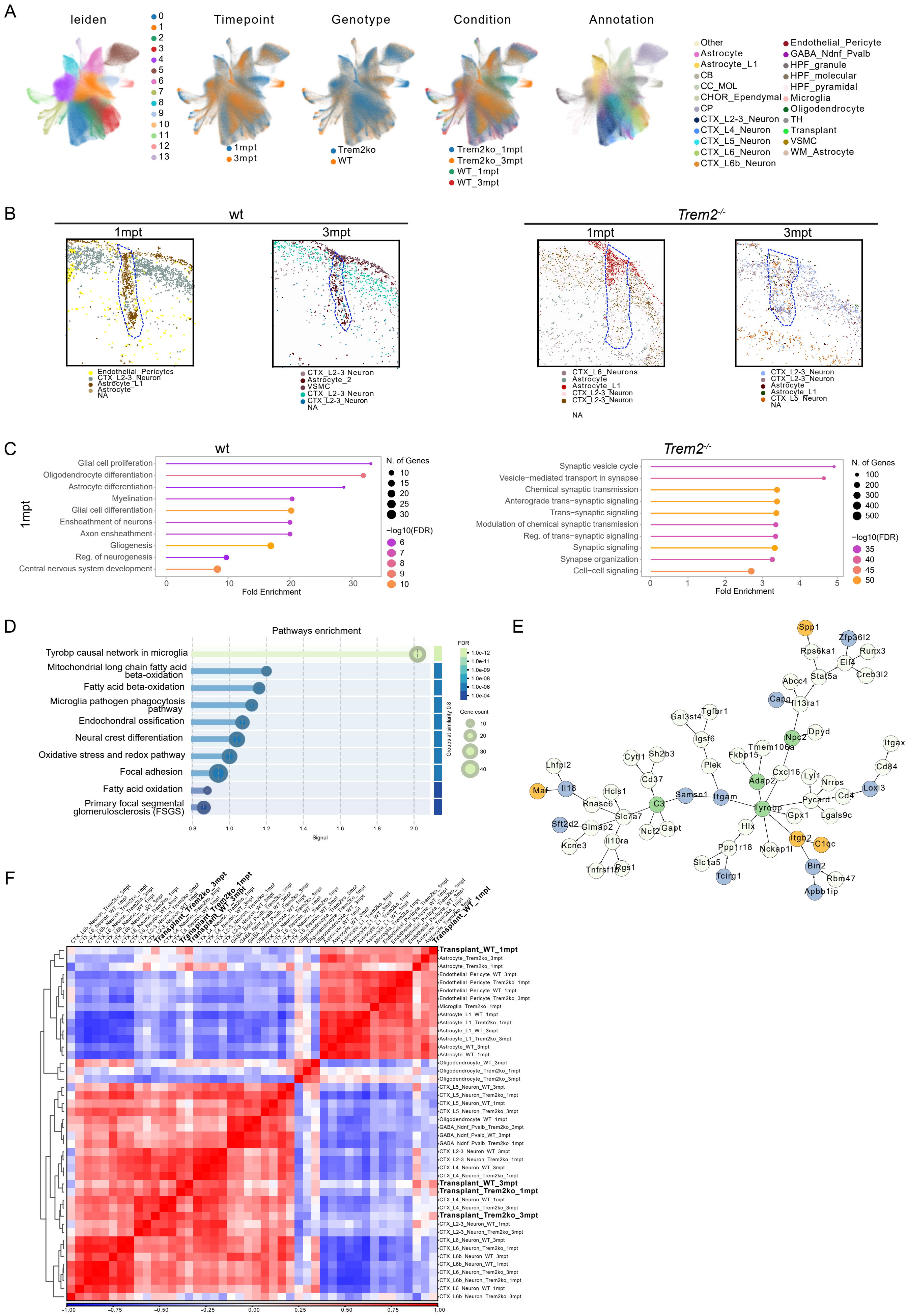
